# Real-time Conformational Dynamics of SARS-CoV-2 Spikes on Virus Particles

**DOI:** 10.1101/2020.09.10.286948

**Authors:** Maolin Lu, Pradeep D. Uchil, Wenwei Li, Desheng Zheng, Daniel S. Terry, Jason Gorman, Wei Shi, Baoshan Zhang, Tongqing Zhou, Shilei Ding, Romain Gasser, Jérémie Prévost, Guillaume Beaudoin-Bussières, Sai Priya Anand, Annemarie Laumaea, Jonathan R. Grover, Lihong Liu, David D. Ho, John R. Mascola, Andrés Finzi, Peter D. Kwong, Scott C. Blanchard, Walther Mothes

## Abstract

SARS-CoV-2 spike (S) mediates entry into cells and is critical for vaccine development against COVID-19. Structural studies have revealed distinct conformations of S, but real-time information that connects these structures, is lacking. Here we apply single-molecule Förster Resonance Energy Transfer (smFRET) imaging to observe conformational dynamics of S on virus particles. Virus-associated S dynamically samples at least four distinct conformational states. In response to hACE2, S opens sequentially into the hACE2-bound S conformation through at least one on-path intermediate. Conformational preferences of convalescent plasma and antibodies suggest mechanisms of neutralization involving either competition with hACE2 for binding to RBD or allosteric interference with conformational changes required for entry. Our findings inform on mechanisms of S recognition and conformations for immunogen design.

---

SARS-CoV-2 spike (S) mediates entry into cells and is a main target for antibody responses against the virus(*1-4*). S is synthesized as a precursor, processed into S1 and S2 by furin proteases, and activated for fusion when human angiotensin-converting enzyme 2 (hACE2) engages the receptor-binding domain (RBD) and when the N-terminus of S2 is proteolytically processed(*5-7*). Structures of soluble ectodomains and native virus particles have revealed distinct conformations of S, including a closed trimer with all RBD oriented downward, trimers with one or two RBDs up, and hACE2-stabilized conformations with up to three RBD oriented up(*6-18*). Real-time information that connects these structures, however, has been lacking. smFRET is well suited to inform on conformational dynamics of proteins reporting domain movements in the millisecond to second range, and has previously been applied to study HIV-1, influenza A, and Ebola spike glycoproteins, via measurements of the distance-dependent energy transfer from an excited donor to a nearby acceptor fluorophores in real-time(*19-23*).

To probe dynamics of SARS-CoV-2 spikes, we used available high-resolution structures of the SARS-CoV-2 S trimer to identify sites of fluorophore pair labeling that have the potential to inform on distance changes expected to accompany conformational changes between the RBD-down and receptor hACE2-induced RBD-up trimer structures(*14, 17*) (Fig. S1). Accordingly, we engineered A4 and Q3 labeling peptides before and after the receptor-binding motif (RBM) to allow site-specific introduction of donor and acceptor fluorophores at these positions (Fig. 1, A, B, and Fig. S1). We optimized retroviral and lentiviral pseudoviral particles carrying the SARS-CoV-2 S protein (Fig. S2) to test the impact of these peptides on infectivity, and found that they were well tolerated, both individually and in combination (Fig. S1D). To measure conformational dynamics of the SARS-CoV-2 S trimer on the surface of virus particles, we established two types of particles, lentiviral pseudoparticles carrying S, as well as coronavirus-like particles generated by expression of S, membrane (M), envelope (E) and nucleocapsid (N) protein (S-MEN)(*24, 25*) (Fig. 1, A and B). S-MEN particles co-express coronavirus surface proteins M and E. Particle quality and the presence of the corona-like S proteins on both particle surfaces were confirmed by cryo-electron microscopy (Fig. 1, C and D).

**Fig. 1.**
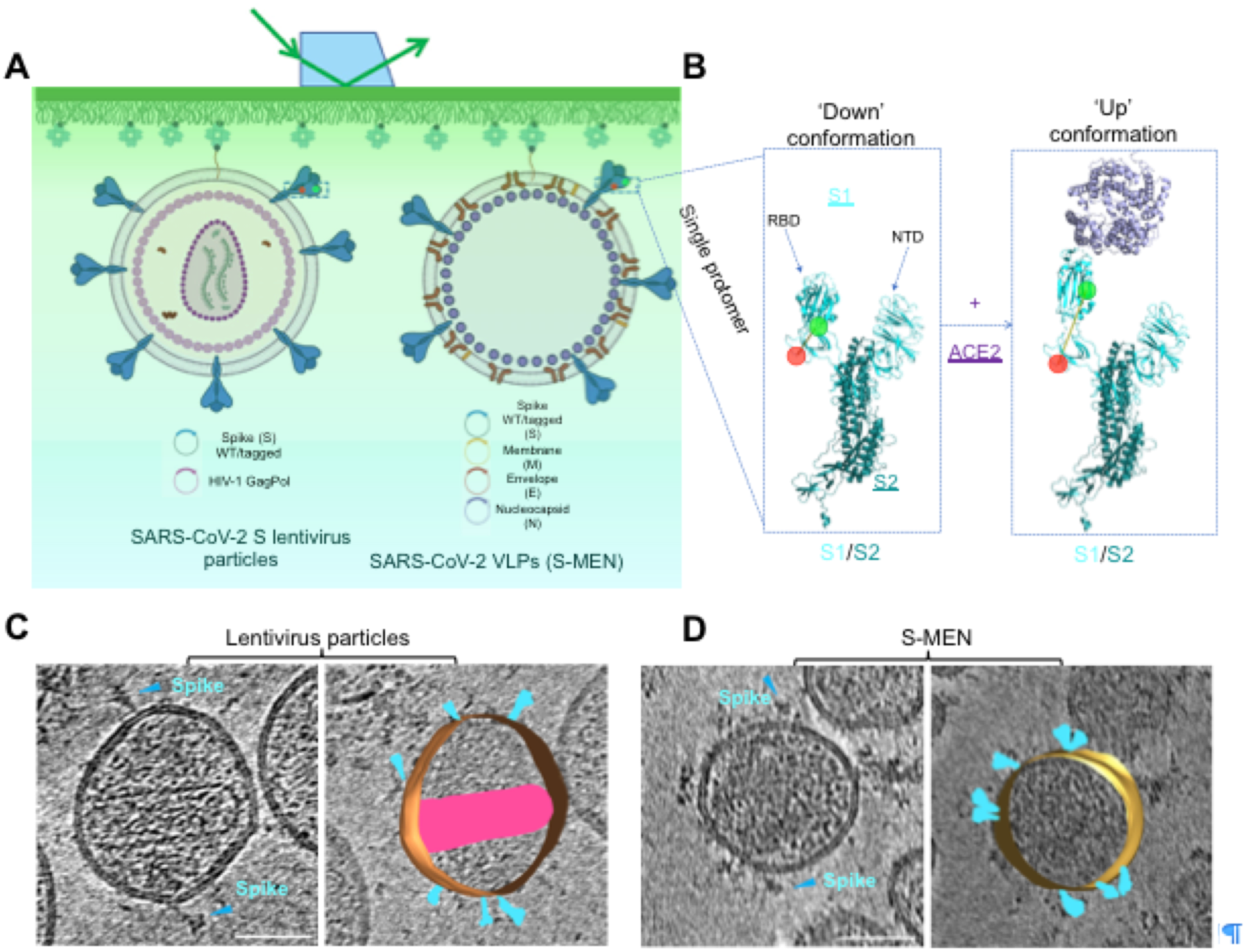
Experimental design for single-molecule FRET imaging of real-time conformational dynamics of the SARS-CoV-2 spike protein on virus particles. **(A, B)** Experimental set-up. **(A)** Virus particles carrying a two-dye labeled SARS-CoV-2 spike protein protomer among wildtype spikes were immobilized on a quartz slide and imaged on a customized prism-based TIRF microscope. Two virus particle systems were developed to incorporate SARS-CoV-2 spike proteins on their surfaces. HIV-1 lentivirus particles comprise HIV-1 cores and SARS-CoV-2 spike proteins on the surface. S-MEN virus-like particles consist of four structural proteins, in which S represents spike, M the membrane protein, E the envelope protein, and N the nucleocapsid protein of SARS-CoV-2. **(B)** Placement of labeling tags for the site-directed introduction of fluorophores (Cy3B, green; LD650, red) was guided by conformational changes in S1 induced by binding of the cellular receptor human angiotensin-converting enzyme 2 (hACE2) from the “RBD-down” to the “RBD-up” conformation (**Fig. S1**). RBD, receptor-binding domain; NTD, N-terminal domain. Structures were adapted from RCSB Protein Data Bank accessories 6VSB (‘Down’ S1/S2 protomer: S1, light cyan; S2, dark blue) and 6VYB/6M0J (‘Up’ protomer S1/S2 engaged with hACE2: hACE2, magenta). **(C, D)** A representative tomographic slice (left panel), and the segmented 3D representation (right panel) of HIV-1 lentivirus particles carrying S (**C**) and S-MEN corona viral-like particles (**D**). Spikes in cyan; HIV-1 capsid in pink. The scale bar is 50 nm.

For smFRET, lentivirus particles and S-MEN particles were generated (see **Materials and Methods**) by transfecting HEK293T cells with an excess of plasmid-encoding wild-type, doped with trace amounts of plasmid expressing labeling peptide-carrying S to ensure the production of virus particles that contain, on average, only a single engineered S protein. As for analogous investigations of HIV-1 Envelope protein(*19, 21*), donor (Cy3B(3S)) and acceptor fluorophore (LD650) were enzymatically conjugated to the engineered S proteins presented on the virus particle surface in situ (see **Materials and Methods**). A biotinylated lipid was then incorporated into the virus particle membrane to allow their immobilization within passivated microfluidic devices coated with streptavidin to enable imaging by total internal reflection microscopy (Fig.1A). Donor fluorophores on single, immobilized virus particles were excited by a single-frequency 532 nm laser and fluorescence emission from both donor and acceptor fluorophores were recorded at 25 Hz (Fig. 2A). From the recorded movies, we computationally extracted hundreds of smFRET traces exhibiting anti-correlated donor and acceptor fluorescence intensities, the telltale signature of conformational changes within the S protein on individual virus particles.

**Fig. 2.**
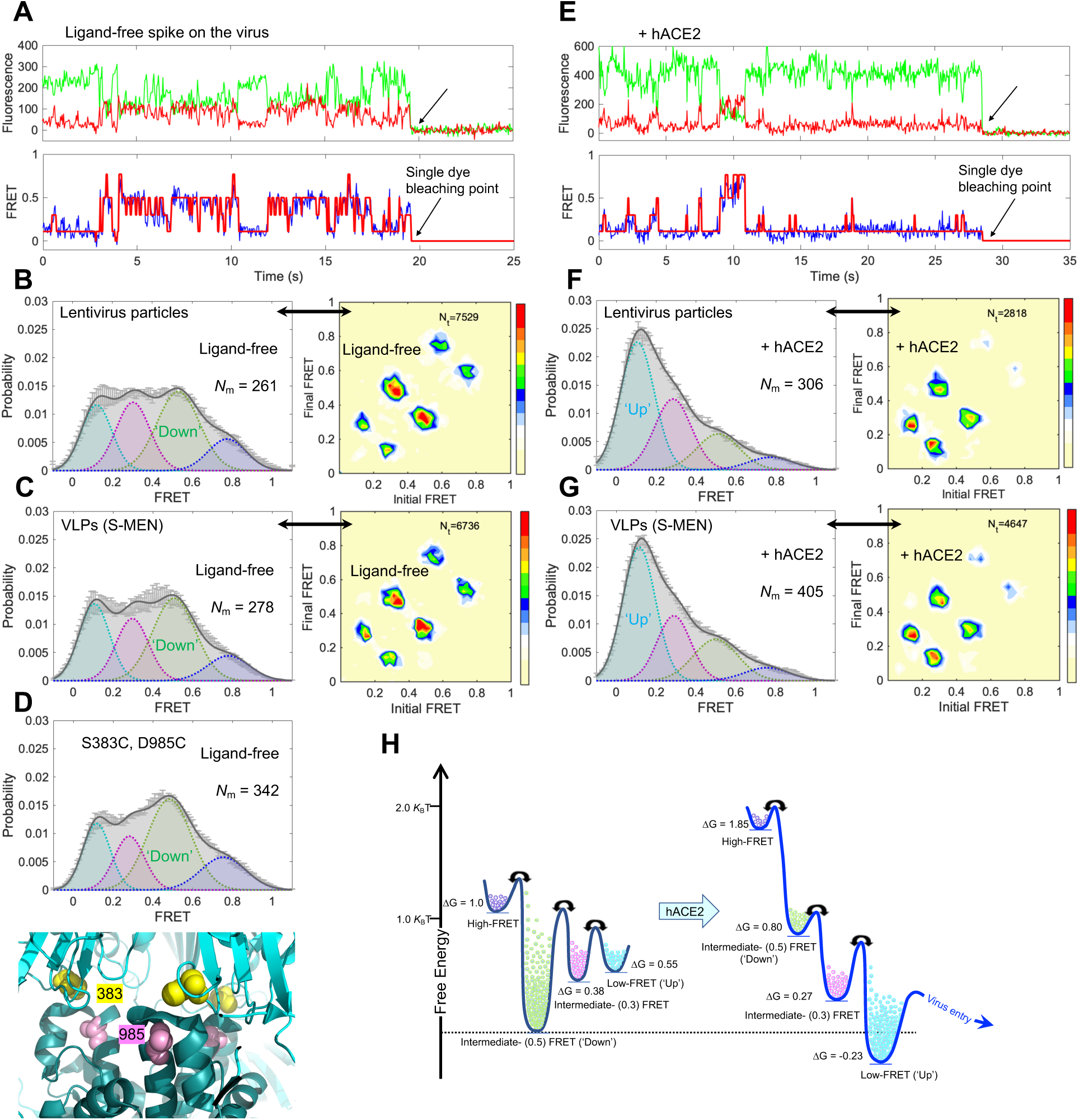
SARS-CoV-2 spike protein is dynamic, and hACE2 shifts conformational landscape from the ground state to the receptor-bound state through one necessary intermediate. **(A–D)**, The ligand-free S on virus particles primarily resides in ‘RBD-down’ conformation (ground-state). **(A)** Example fluorescence trace (Cy3B, green; LD650, red) and resulting quantified FRET traces (FRET efficiency, blue; hidden Markov model initialization, red) of a dually labeled ligand-free spike protein on the surface of HIV-1 lentivirus particle. Arrows point to the single-step photobleaching steps of dyes at the single-molecule level and define the baseline. (**B, C)** FRET histograms (left) and transition density plots (TDPs, right) of ligand-free spikes on lentivirus particles (**B**) and S-MEN viral-like particles (**C**). Number (N) of individual dynamic traces compiled into a conformation-population FRET histogram (gray lines) and fitted into a 4-state Gaussian distribution (solid black) centered at 0.1-FRET (dashed cyan), 0.3-FRET (dashed red), 0.5-FRET (dashed green), and 0.8-FRET (dashed magenta). TDPs, displayed as initial FRET vs. final FRET with relative frequencies, trace the locations of state-to-state transitions and their relative frequencies (max red scale = 0.01 transitions/second), originated from the idealization of individual FRET traces in FRET histograms. **(D)** A modified spike (S383C and D985C)(*9, 11*) stabilized in ‘RBD-down’ conformation, observed from the FRET histogram (upper panel). The small increase in the population of the ground state (∼0.5 FRET) likely reflects the partial nature of the formation of the disulfide in this mutant, which has 40% the infectivity of wildtype (**Fig. S3C**). Modified S383C and D985C depicted in the high-resolution structure of S 6ZOY (lower panel). (**E–G)**, Experiments as in **A–C**, respectively, conducted in the presence of 200 μg/ml monomeric hACE2. The soluble hACE2 activates spike proteins on the virus by shaping the conformational landscape towards stabilizing the ‘RBD-up’ conformation (activated-state). FRET histograms represent mean ± s.e.m., determined from three randomly assigned populations of all FRET traces under corresponding experimental conditions. N, number of individual FRET traces. Evaluated state occupancies see **Table S1. (H)** Relative free energy model of conformational landscapes of SARS-CoV-2 spikes in response to the hACE2 binding. The differences in free energies between states were roughly scaled based upon relative state occupancies of each state.

Analyses of smFRET data from ligand-free S protein on lentiviral particles revealed that the SARS-CoV-2 S protein is dynamic, sampling at least four distinct conformational states characterized by low-(∼0.1), intermediate-(∼0.3 and ∼0.5), and high-(∼0.8) FRET efficiency (FRET). Population FRET histograms, comprised of hundreds of smFRET traces, revealed that the conformation exhibiting intermediate FRET (∼0.5) to be the most abundantly occupied (Fig.2B, and Table S1). Comparable findings were obtained for S protein incorporated into S-MEN coronavirus-like particles (Fig. 2C). Notably, the intermediate-FRET (∼0.5) conformation of the S protein was stabilized by a disulfide bridge between amino acids 383 and 985 (Fig. 2D, Fig.S3, A and B). Because electron microscopy (EM) methods have identified the disulfide-bridge stabilized state as the three-RBD-down conformation(*9, 11*), these experiments suggest the intermediate-(∼0.5) FRET S protein conformation to represent the closed trimer with all three RBD oriented downward (RBD-down conformation).

To identify the receptor-bound conformation of the SARS-CoV-2 S protein by smFRET, we measured the conformational consequences of soluble, monomeric hACE2 binding. Addition of the monomeric hACE2 receptor to surface-immobilized virus particles lead to increased occupancy of the low-(∼0.1) FRET S protein conformation (Fig. 2E), which was observed at both the single-molecule and population level (Fig. 2F). Similar hACE2 receptor impacts on the SARS-CoV-2 S protein were observed in both lentiviral particle and S-MEN coronavirus-like particle contexts (Fig. 2, E to G). Dimeric hACE2, a more potent ligand (Fig. S4A)(*26*), stabilized the low-(∼0.1) FRET S protein conformation more efficiently (Fig. S4, B and C), suggesting that the observed low-FRET state likely represents the receptor-bound state in which all three RBD domains are oriented upwards (RBD-up conformation).

A unique strength of single-molecule imaging is its capacity to reveal directly both the structural and kinetic features that define biological function(*27, 28*). To extract such information for the SARS-CoV-2 S protein, we employed Hidden Markov Modeling (HMM)(*29*) to idealize individual smFRET traces. These data allowed quantitative analyses of the thousands of discrete FRET transitions observed for the S protein on the surface of lentiviral and S-MEN particles in the absence and presence of ligands to gain insights into the order and timing of conformational changes across conditions (Figs. 2, B, C, F, G, and Fig. S4). Transition density plots (TDPs), which conveniently display the nature, order and frequency of the transitions observed in the ensemble of molecules examined(*27*), immediately revealed two salient features that informed on the nature of S protein dynamics. First, the symmetry of the evidenced transitions in each TDP with respect to the diagonal axis indicated that the unliganded S protein is in equilibrium exchange between distinct conformations under ambient conditions - i.e. physiological buffer at room temperature. Second, the transitions evidenced in the TDP exhibited a defined transition order, from low- to intermediate-FRET and from intermediate- to high-FRET conformations. These findings suggest that the SARS-CoV-2 S protein exhibits a defined sequence of activating structural transitions wherein the ground state RBD-down conformation (intermediate-(∼0.5) FRET) converts to the receptor-activated RBD-up conformation (low-(∼0.1) FRET) via at least one intermediate-(∼0.3) FRET conformation. They further reveal that the RBD-down conformation of the S protein can reversibly access at least one additional high-(∼0.8) FRET conformation. The most frequent transitions evidenced were those between RBD down (∼0.5 FRET) and the on-path intermediate-(∼0.3) FRET states, from which relatively infrequent transitions to the low-FRET (∼0.1), RBD-up conformation - akin to those observed upon hACE2-binding – could be achieved spontaneously.

As expected, the binding of the hACE2 receptor modified the dynamic S protein conformational landscape towards the RBD-up conformation (∼0.1 FRET), rendering it the most populated (Fig. 2, B, C, F, G). This change resulted from an increase in the transition rate from the RBD-down conformation (∼0.5 FRET) towards the intermediate-(∼0.3) FRET state and subsequently the RBD-up (∼0.1 FRET) conformation, which was also modestly stabilized. The energy barriers for reverse transitions towards the RBD-down conformation (∼0.5 FRET), were also elevated, explaining receptor-bound conformation accumulation over time (Figs. S5). These analyses lead to a qualitative model for hACE2 activation of the SARS-CoV-2 S protein from the ground state to the receptor-bound state through at least one intermediate conformation (Fig. 2H). The summary of relative state occupancies, transition rates among conformations and errors are listed in Tables S1 and S2, respectively.

In most cell types, the serine protease TMPRSS2 is required for pH-independent SARS-CoV-2 entry(*5, 30, 31*). In vitro, the effect of TMPRSS2 is mimicked by the serine protease trypsin, which has similar cleavage specificity(*5, 31*). smFRET analysis of trypsin-treated S protein on lentiviral particles in the absence of receptor revealed a clear shift towards activation (Fig. 3, A, B). After trypsin treatment, the addition of hACE2 receptor was more effective at stabilizing the S protein in the RBD-up (∼0.1 FRET) conformation (Fig. 3, C and D, Fig. S6). To further validate this finding, we measured the effects of trypsin pre-treatment in virus-cell and cell-cell fusion assays using split NanoLuc system consisting of LgBiT and HiBiT (Fig. 3, E and F, Fig. S7). Here, membrane fusion restored luciferase activity between lentiviral particles carrying the S protein as well as a Vpr-HiBiT fusion protein with cells expressing the LgBiT counterpart fused to a PH domain. This assay revealed fusion to be strictly dependent on the hACE2 receptor and to be stimulated by trypsin treatment (Fig. 3, E and F). Nearly identical results were observed for cell-cell fusion between donor cells expressing S and target cells expressing hACE2 (Fig. S7), confirming the activating role of trypsin treatment.

**Fig. 3.**
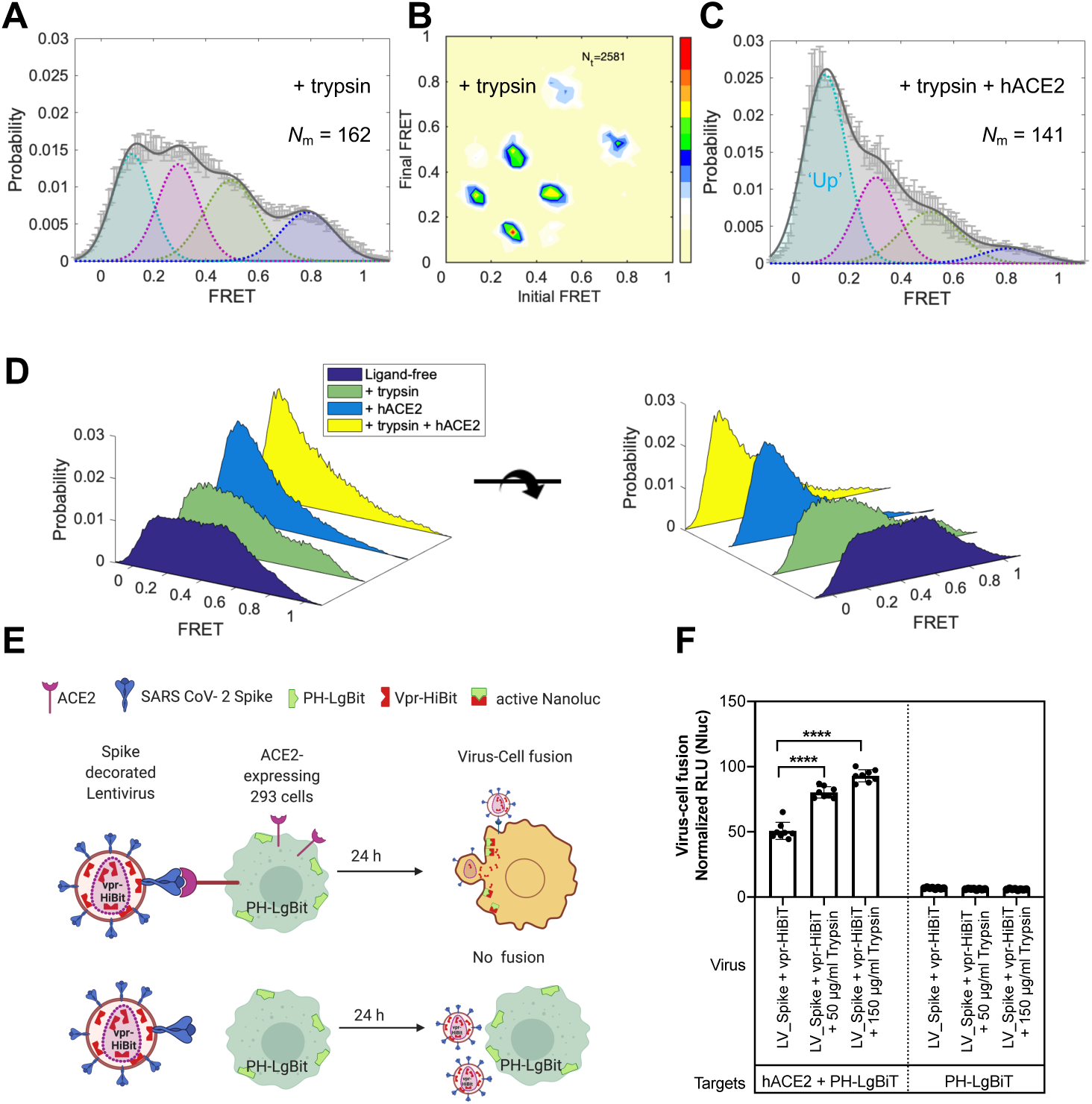
Conformational effects of trypsin treatment of spikes follow the hACE2-dependent activation pathway. **(A–C)** The serine protease trypsin remodels conformational landscape of spike proteins towards down-stream conformations on the path of hACE2-dependent activation. **(A, B)** The FRET histogram (**A**) and TDP (**B**) of spike proteins on HIV-1 lentivirus particles in the presence of 50 μg/ml trypsin. **(C)**, An experiment as in **(A)**, for spikes in the presence of both 50 μg/ml trypsin and 200 μg/ml hACE2. **(D)** Three-dimensional presentations of FRET histograms of spike proteins on the virus in the presence and the absence of hACE2 and trypsin. FRET histograms represent mean ± s.e.m., determined from three randomly assigned populations of FRET traces. Evaluated state occupancies see **Table S1. (E, F)** Trypsin enhances SARS-CoV-2 spike-mediated hACE2-dependent virus-cell fusion. **(E)** Assay design to monitor virus-cell fusion using the HiBit and LgBiT split NanoLuc system(*47*). Vpr-HiBit was packaged into lentiviral particles carrying SARS-CoV-2 spike. HEK293 target cells transiently expressing LgBiT tagged to PH domain of human phospholipase Cd at the N-terminus alone or together with hACE2. hACE2-dependent virus-cell fusion was determined by monitoring reconstituted NanoLuc activity in target cells 24 h after infection. **(F)** Normalized relative luciferase units (RLU; mean ± s.d., two replicates with quadruplicates) measured 24 h post infection to quantify virus-cell fusion in stated target cells after treating viruses with or with indicated amounts of trypsin for 15-20 min at 37 °C. NanoLuc activities were normalized to luciferase activity detected in uninfected target cells. p values derived from unpaired t-test; **** corresponds to p < 0.0001.

We next explored the suitability of the smFRET assay to characterize the conformational consequences of antibody binding to the SARS-CoV-2 S protein. Multiple studies on antibodies generated from COVID-19 patients have shown that one type of antibody often dominates immune responses(*32-37*). This prompted us to screen plasma from convalescent patients with neutralizing activity that can bind to the S protein on lentiviral particles(*38*) using a modified virus-capture assay (VCA)(*39*). Cross-reactive CR3022(*40*), one of the very first reported antibodies from SARS-CoV-1 that also bind to SARS-CoV-2 spike RBD domain, served as a good indicator of RBD binding (Fig. 4A). We identified two plasma samples (S002 and S006) able to specifically bind the RBD, recognize S expressed at the cell surface and to neutralize viral particles (Fig. 4, A to C, and Fig. S8). smFRET analysis of antibody-bound S revealed that both CR3022 and plasma from patient S006, stabilized S in the RBD-up (∼0.1 FRET) conformation, in a similar fashion as receptor hACE2 (Fig. 4, D and E). These data point to the presence of RBD-directed antibodies in patient S006. By contrast, smFRET indicated that plasma from patient S002 contained an activity that stabilized the RBD-down (∼0.5 FRET) conformation (Fig. 4F). Plasma S002 antagonized hACE2 binding, but RBD competition did not affect its recognition of S, suggesting that its neutralization activity does not solely rely on blocking the receptor interface We then assessed the conformational preference of four RBD-directed antibodies: the potently neutralizing antibodies H4, 2-4 and 2-43, and the neutralization nanobody VHH72, each of which binds RBD in a different way(*41-43*). Antibody H4 and nanobody VHH72 stabilized the S protein in an RBD-up (∼0.1 FRET) conformation similar to hACE2, CR3022, and S006, whereas antibody 2-4 shifted the conformational landscape towards RBD-down (∼0.5 FRET) conformation, similar to S002 (Fig. 4, G to J). The very potent neutralizing antibody 2-43(*43*), meanwhile, showed a partial shift to the RBD-up (∼0.1 FRET) conformation (Fig. S9). The absence or presence of hACE2 did not appear to affect the RBD-up stabilization evidenced for antibodies CR3022, S006, VHH72, or H4 (Fig. S9). However, plasma S002, and to a lesser extent antibody 2-4, reduced the hACE2-dependent stabilization of the RBD-up (∼0.1 FRET) conformation, suggesting that they may interfere with hACE2 receptor binding via an allosteric mechanism. These findings indicate that SARS-CoV-2 neutralization can be achieved in two ways: 1) antibodies that conformationally mimic hACE2 and directly compete with hACE2 receptor binding, or 2) by allosterically stabilizing the S protein in its RBD-down conformation.

**Fig. 4.**
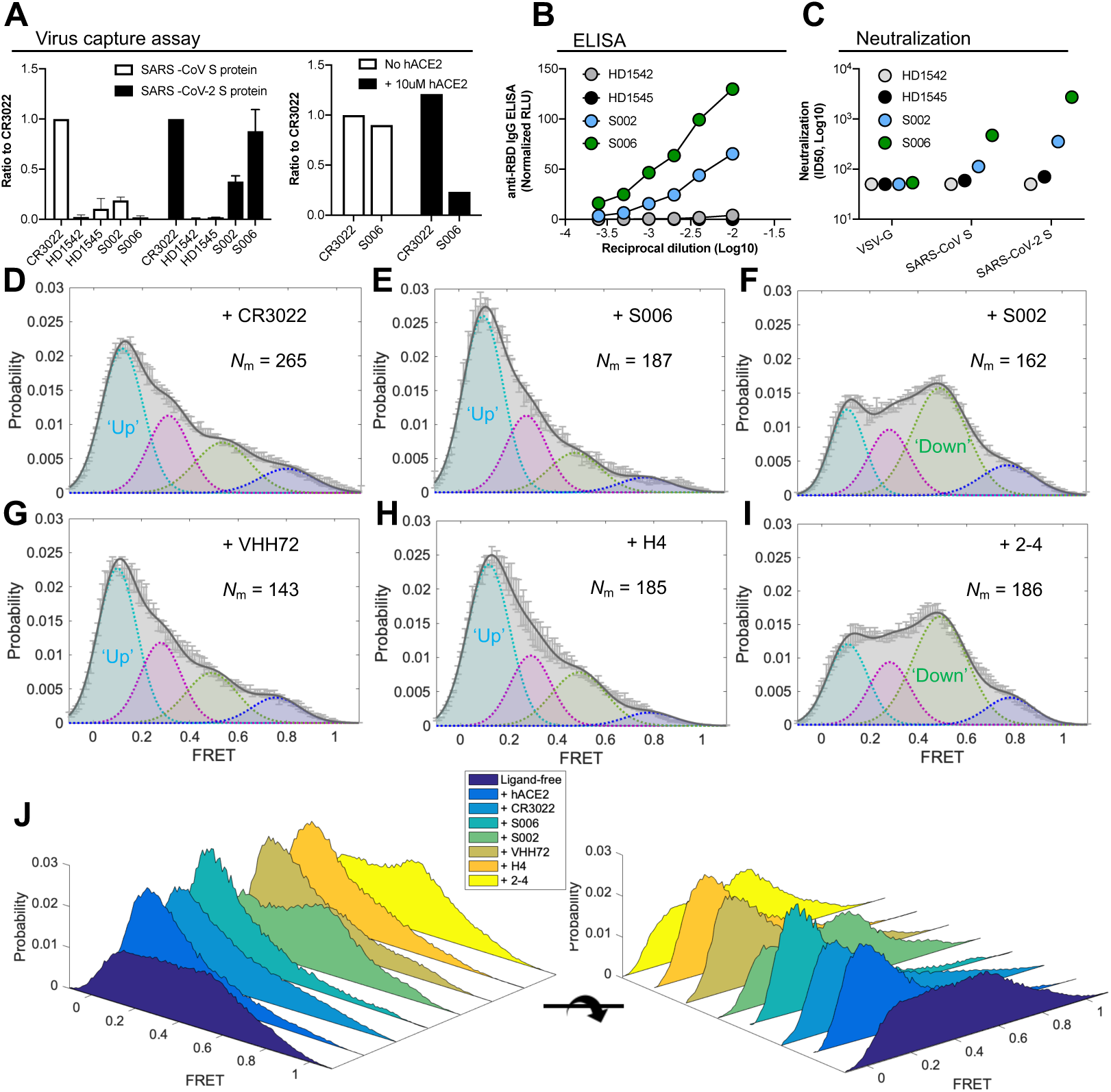
Spike-conformational preferences of RBD-directed monoclonal antibodies and convalescent patient plasma. **(A)** Bar graphs of a virus capture assay showing that convalescent plasma from SARS-CoV-2-positive patients S006 and S002 can bind to SARS-CoV-2 S on viral particles not to SARS-CoV-1 in reference to a cross-reactive SARS-CoV-1 monoclonal antibody CR3022. Plasma from healthy donors HD1542 and HD1545 as controls. Unless noticed, S006, S002, HD1542, and HD1542 represent plasma. The right panel depicts sensitivity of S006 binding to S to soluble ACE2. Virus capture experiments were repeated three times and represented as mean ± s.d. **(B)** The binding affinity of both S006 and S002 towards SARS-CoV-2 RBD in comparison to plasma from heathy donors by anti-RBD IgG ELISA. **(C)** Both S006 and S002 show virus neutralizing ability against lentiviruses carrying SARS-CoV-2 S. **(D–I)** FRET histograms of CR3022 (**D**) at 200 μg/ml, convalescent patient plasma from S006 (**E**) and S002 (**F**) at 1:100 dilution, and RBD-directed monoclonal antibodies VHH72 (**G**), H4 (**H**), and 2-4 (**I**) at 200 μg/ml. **(J)** Three-dimensional presentations of FRET histograms of spikes on the retrovirus under different conditions (**D–I**) in reference to ligand-free (**Fig. 2B**). FRET histograms represent mean ± s.e.m., determined from three randomly assigned populations of FRET traces. For state occupancies see **Table S1**.

The strength of the presented smFRET approach is revealed by the capacity to examine the dynamic properties of the S protein in real time, including: 1) the distinct conformational states that it spontaneously transits under physiological conditions; 2) the impact of sequence alterations on S protein dynamics; and 3) the responses of the S protein to cognate hACE2 receptor and antibody recognition.

The present analyses of dynamic S protein molecules provides three lines of evidence that indicate that the intermediate-(∼0.5) FRET state observed represents the RBD-down, ground state conformation of the S protein, in which all three RBD domains are oriented towards the viral particle membrane. First, in line with previous electron microscopy (EM) investigations(*6-16*), the RBD-down state is the most populated. In further agreement with recent EM studies, both the disulfide bridge (S383C, D985C)(*9, 11*) and antibody 2-4 stabilized the S protein in a conformation with all three RBD oriented down^42^. While our smFRET observations highlight considerable conformational flexibility in these contexts compared to EM of soluble trimers, we attribute these distinctions to a tendency of our analysis approach to over-emphasize dynamic features, while EM may over-emphasize static conformations rigidified by cryogenic temperatures that may be more readily identified and classified(*44*). Multiple lines of evidence also facilitated assignment of the RBD-up (∼0.1 FRET) conformation of the S protein with all three RBD domains oriented away from the virus particle membrane. For instance, this conformation was stabilized by soluble monomeric hACE2 receptor, and even further stabilized in the presence of soluble, dimeric hACE2 receptor as well as RBD-targeting antibodies, such as CR3022, that are known to access their epitopes when the S protein is in an activated, RBD-up conformation(*40-42*). The structure of the on-path (∼0.3 FRET) intermediate observed during S1 opening is likely similar to the all-down ground state; cryo-EM structures of soluble SARS-CoV-2 S trimers(*17*) that engage one or two hACE2 molecules receptors(*18*) reveal that the distance between the two labeling sites increases in the ligand-free protomers adjacent to a protomer engaged to hACE2 (Fig. S1A). The additional, highly compacted S conformation (∼0.8 FRET) evidenced, which is also depopulated by activating ligands, remains unknown.

These smFRET analyses are in global agreement with the conformational states observed at the single particle EM and cryoET level(*6, 8, 9, 11-17, 40-42, 45, 46*). The observed FRET changes are also are in good agreement with expected increase in the distance between the labeling peptide insertion sites that carry the fluorophores in the RBD-down and RBD-up conformations of the S trimer. The capacity to examine the conformational preferences of RBD-directed antibodies to the S protein enabled us to identify conformational signatures of antibodies in patient plasma. This approach identified patients with antibody activities that either mimicked ACE2 (indicating anti-RBD activity) or stabilized the ground state of S, thereby interfering with hACE2 receptor-mediated activation of the SARS-CoV-2 S protein. We anticipate that ground state stabilization by antibodies, small molecules, or the rational engineering of trimers may represent an effective avenue of antagonism as well as immunogen design.

## Acknowledgments

We thank Dr. Craig Wilen for sharing Huh7.5 cells, Dr. Nevan Krogan for sharing M, E and N expression plasmids, Dr. Marc Johnson for sharing pCMV-S Δ19 and HIV-1 GagPol-InGluc plasmids, and MLV In-GLuc and Dr. David Derse for sharing HIV In-GLuc plasmids. This work was supported by NIAID, NIH grant R01 AI150560 to W.M. and S.C.B, by NIGMS, NIH R01 GM098859 to S.C.B., by the Intramural Research Program of the Vaccine Research Center, NIAID, NIH to P.D.K., and by Ministère de l’Économie et de l’Innovation du Québec, Programme de soutien aux organismes de recherche et d’innovation and by the Fondation du CHUM to A.F. A.F. is the recipient of a Canada Research Chair on Retroviral Entry # RCHS0235 950-232424. R.G. is supported by a MITACS Accélération postdoctoral fellowship. J.P., G.B.B. and S.P.A. are supported by CIHR graduate fellowships.

## Author contributions

M.L. and W.M. designed the studies. M.L. performed mutagenesis, virus infectivity assays, generated fluorescently labeled viruses, performed smFRET imaging and analyzed the data. P.D.U. designed and performed virus-cell fusion, cell-cell fusion and sACE2 inhibition assays. W.L. generated tomograms of virus particles. W.L., J.R.G. and P.D.U. assisted with virus-like particles. D.Z. built the prism-TIRF microscope with assistance from M.L. D.S.T and S.C.B, who supported and advised on the development and implementation of single-molecule instrumentation, data acquisition and analysis in the Mothes lab. B.Z., T.Z., J.G., L.L., D.D.H, and P.D.K. provided various antibodies, and scientific inputs into experiments and manuscript editing. W.S. and J.R.M. provided monomeric and dimeric hACE2. A.L. and A.F. processed all the blood sample and prepared patient plasma. S.D. performed virus capture assay. R.G., S.D., and A.L., performed neutralization experiments. J.P. and S.P.A. performed flow cytometry experiments. M.L. and W.M. wrote the manuscript with input from all authors.

## Data and materials availability

All data is available in the main text or the supplementary materials. The data that support the findings of this study are available from the corresponding authors upon reasonable request. The full source code of SPARTAN, which was used for analysis of smFRET data, is publicly available. (http://www.scottcblanchardlab.com/software). Some small customized Matlab scripts are available upon reasonable requests.

## Materials and Methods

### Construction of full-length tagged SARS-CoV-2 spike (S)

A full-length wild-type pCMV3-SARS-CoV-2 Spike (S1+S2)-long (termed as pCMV-S, codon-optimized, Sino Biological, cat # VG40589-UT) plasmid was used as a template to generate tagged pCMV-S. The translated amino acid sequence of pCMV-S is identical to QHD43416.1 (GenBank). Labeling tags Q3 (GQQQLG) and A4 (DSLDMLEM)(*48, 49*) were placed before and after the receptor-binding motif (RBM) where the cellular receptor ACE binds (Fig. S1). The short peptide labeling tag Q3 was inserted at either of two positions in the receptor-binding domain (RBD) to generate the pCMV-S Q3-1 and Q3-2 constructs. The labeling tag A4 was inserted at either of two sites in subunit domain 1 (SD1) to generate the constructs pCMV-S A4-1 and A4-2, respectively. Insertion of single tags into S proteins did not affect viral infectivity. Four different combinations of double-tagged S (designated as pCMV-S Q3-1 A4-1, Q3-1 A4-2, Q3-2 A4-1, Q3-2 A4-2) were constructed to permit smFRET imaging. Each pair of inserted tags did not compromise S-dependent lentivirus infectivity.

### Infectivity measurements

The infectivity of lentivirus particles carrying SARS-CoV-2 spike proteins was determined using a vector containing an HIV-1 or MLV long terminal repeat (LTR) that expresses a Gaussia luciferase reporter (Gluc)(*50, 51*) 293T cells were cultured in DMEM (Gibco) media, supplemented with 10% FBS (Gibco), 100 U/ml penicillin/streptomycin (Gibco), 2 mM L-glutamine (Gibco), and in the presence of 5% CO_2_. Cell culture media was exchanged before transfection. Cells were transfected at 60–80 % confluency with the plasmid encoding respective spike protein, the plasmid encoding an intron-regulated Gluc (HIV-1-inGluc), and a plasmid pCMV delta R8.2 encoding HIV-1 GagPol (Addgene, plasmid #12263) using Fugene 6 (Promega, # E2311). Variants of plasmids encoding spike included the wild-type pCMV-S, single or dually tagged pCMV-S, pCMV-S Δ19 (S lacking the last 19 amino acids of the cytoplasmic tail), and an additional S Δ19 expressing plasmid (a gift from Mark Johnson, University of Missouri). To generate retroviral particles with an MLV core carrying full-length S, an MLV GagPol construct, and an intron-regulated Gaussia luciferase reporter construct (MLV-inGluc) were used(*50*). An HIV-1 GagPol-InGluc construct(*52*) (a gift from Mark Johnson, University of Missouri) in which HIV-1 GagPol and HIV-1-inGluc genes are expressed from a single plasmid were also tested. Viruses were harvested 40 hours post-transfection, filtered, and concentrated 7-fold by ultracentrifugation at 25,000 rpm in an SW28 rotor for 2 hours at 4 °C. Pelleted viruses were resuspended in above DMEM media and titered on Huh7.5 cells that endogenously express hACE2. Gluc activity was measured in the cell supernatant at 48 hours using a Pierce™ Gaussia Luciferase Flash Assay Kits (ThermoFisher Scientific, # 16158).

### Soluble human ACE2 (hACE2) inhibition assay

NanoLuc-expressing SARS-CoV-2 decorated lentiviruses were first generated by transfecting a plasmid mixture containing 5 μg of pCMV-S, 5 μg of pCMV delta R8.2 (expresses HIV-1 GagPol), 2 μg of pLL3.7 NanoLuc (packaging reporter to express NanoLuc) into 60-70 % HEK293 cells using FuGENE 6 (3ul of FuGENE 6 for every µg of DNA) in a 10 cm TC dish. The culture supernatants were harvested at 48 h after transfection, clarified using 0.45 µm filter, and viruses partially purified by sedimentation through a layer of 15 % sucrose solution made in 1X PBS. The virus pellet was resuspended in OptiMEM to achieve a 20-fold concentration over original culture volume. Infectivity inhibition assays were performed by incubating NanoLuc expressing lentiviruses either alone or with soluble hACE2 (monomeric and dimeric) at indicated concentrations in triplicates for 90 mins at room temperature prior to infection of the 293T cells stably expressing hACE2 seeded in a tissue-culture treated 96-well solid white assay plate (Costar Inc, catalog #3917). The culture supernatants were completely removed 24 h p.i. and cells lysed using 50 µl NanoLuc assay buffer. Luciferase activity was then measured using Tristar multiwell luminometer (Berthold Technology, Bad Wildbad, Germany) for 2.5 seconds by adding 20 µl of Nano-Glo® substrate (Promega Inc, WI, USA; diluted 1:40 in NanoLuc assay buffer). HEK293T cells (not expressing hACE2) infected similarly served as controls to determine basal luciferase activity for obtaining normalized relative light units in infected samples. Infectivity obtained in mock-treated samples were set to 100 %. The data were processed and plotted using GraphPad Prism 8 v8.4.3.

### Preparations of convalescent plasma from SARS-CoV-2-positive patients

All SARS-CoV-2-postive patient serum work was conducted in accordance with the Declaration of Helsinki in terms of informed consent and approval by an appropriate institutional board (CHUM, 19.381). The donors met all donor eligibility criteria: previous confirmed COVID-19 infection by PCR and complete resolution of symptoms for at least 14 days.

Donor S002: Male, 65 years old, sample recovered 25 days after symptoms onset.

Donor S006: Male, 30 years old, sample recovered 41 days after symptoms onset.

### Antibody neutralization assays

In the case of VSVG, SARS-CoV-1, and SARS-CoV-2 parallel neutralization assay (Fig. 4C, and Fig. S8C), experiments were performed as follows. Target cells were infected with single-round luciferase-expressing lentiviral particles. Briefly, 293T cells were transfected by the calcium phosphate method with the pNL4.3 R-E-Luc plasmid (NIH AIDS Reagent Program) and a plasmid encoding for SARS-CoV-2 Spike, SARS-CoV Spike or VSV-G at a ratio of 5:4. Two days post-transfection, cell supernatants were harvested and stored at –80 °C until use. 293T-ACE2(*53*) target cells were seeded at a density of 1×10^4^ cells/well in 96-well luminometer-compatible tissue culture plates (Perkin Elmer) 24 hours before infection.

Recombinant viruses in a final volume of 100 μl were incubated with the indicated sera dilutions (1/50; 1/250; 1/1250; 1/6250; 1/31250) for 1hours at 37°C and were then added to the target cells followed by incubation for 48 hours at 37°C; cells were lysed by the addition of 30 μl of passive lysis buffer (Promega) followed by one freeze-thaw cycle. An LB941 TriStar luminometer (Berthold Technologies) was used to measure the luciferase activity of each well after the addition of 100 μl of luciferin buffer (15 mM MgSO_4_, 15 mM KH_2_PO_4_ [pH 7.8], 1 mM ATP, and 1 mM dithiothreitol) and 50 μl of 1 mM d-luciferin potassium salt (Prolume). The neutralization half-maximal inhibitory dilution (ID_50_) represents the plasma dilution to inhibit 50 % of the infection of 293T-ACE2 cells by recombinant viruses bearing the indicated surface glycoproteins. The plasmids expressing the human coronavirus Spikes of SARS-CoV-2, SARS-CoV, NL63 229E and OC43 were previously described(*38, 53*). The pNL4.3 R-E-Luc was obtained from NIH AIDS Reagent Program. The plasmid encoding for SARS-CoV-2 S RBD was reported elsewhere(*53*). The vesicular stomatitis virus G (VSV-G)-encoding plasmid (pSVCMV-IN-VSV-G) was previously described(*53*).

### Virus-cell fusion assays

We used the split nanoluc assay to monitor virus-cell fusion(*47*). For the preparation of Vpr-HiBiT–containing SARS CoV-2 decorated lentiviruses, a plasmid mixture containing 4 μg of pCMV-S, 4 μg of psPAX2 (Gag-pol, Rev, and Tat expression vector; does not express Vpr), 2 μg of pLL3.7 (packaging reporter to express GFP), and 2 μg of pCMV-Vpr-HiBiT (Vpr-HiBiT expression vector)) was transfected into 60-70 % HEK293 cells using FuGENE 6 (3ul of FuGENE 6 for 1µg of DNA). The culture supernatants were harvested at 48 hours after transfection, clarified using 0.45 µm filter, and viruses partially purified by sedimentation through a layer of 15 % sucrose solution made in 1X PBS. The virus pellet was resuspended in OptiMEM to achieve a 20-fold concentration over original culture volume. Virus preparations were mock-treated or treated with trypsin (50 and 150 µg/ml) at 37 °C for 15 min. Neat serum at a final concentration of 10 % was added to terminate both trypsin and mock-treated samples. 2.5 × 10^5^ HEK293 cells in 24-wells were transfected with 400 ng of hACE2-expressing plasmid or pcDNA3.1 vector along with 100 ng PH-LgBiT (LgBiT-tagged to pleckstrin homology domain of human phospholipase Cd at the N-terminus) expressing plasmid using FuGENE 6 for use as target cells. 24 hours post-transfection, 2 × 10^5^ cells were seeded in tissue-culture treated 96-well solid white assay plate (Costar Inc, #3917) in 100 µl RPMI complete medium. They were infected with 25 μl of prepared Vpr-HiBiT-containing viruses in quadruplicates for 24 hours at 37 °C. Light signals derived from reconstituted HiBiT/LgBiT association were measured using Tristar multiwell luminometer (Berthold Technology, Bad Wildbad, Germany) for 2.5 seconds by adding 20 µl of Nano-Glo® substrate (Promega Inc, WI, USA; diluted 1:40 in PBS). Mock-infected cells served as controls to determine basal luciferase activity for obtaining normalized relative light units in infected samples. The data were processed and plotted using GraphPad Prism 8 v8.4.3.

### Cell-cell fusion assays

2.5 × 10^5^ HEK293 cells seeded per well of 24-well plate were transfected with 400 ng of hACE2 and 100 ng PH-LgBiT expressing plasmid(*47*) using FuGENE 6 (3 ul of FuGENE 6 for 1 µg of DNA). The second set of HEK293 cells for co-culture were transfected with 400 ng of SARS CoV-2 spike-expressing plasmid or pcDNA 3.1 vector along with 100 ng of vpr-HiBiT (HiBiT tagged to HIV-1 Vpr at the N-terminus) expressing plasmid. 24 hours later, HEK293 cells expressing Vpr-HiBiT alone or co-expressing spike and Vpr-HiBiT were either untreated or treated with 50 µg/ml trypsin solution for 10 min at 37 °C. Trypsin was neutralized by adding serum-containing medium and cells washed once before resuspending in complete RPMI medium. 1 × 10^5^ cells from various Vpr-HiBiT and PH-LgBiT expressing populations were mixed (1:1 ratio, 50 µl each population) in tissue culture treated 96-well solid white assay plate (Costar Inc, catalog #3917) in quadruplicates. Membrane fusion was initiated by spinning down the mixed cells in a 96-well plate (1600 rpm for 5 min, swing-out buckets, Heraeus, Sorvall centrifuge). The cells were then incubated at 37 °C in tissue culture incubator for 4 hours or 24 hours before measuring reconstituted HiBiT/LgBiT activity using a Tristar multiwell Luminometer (Berthold Technology, Bad Wildbad, Germany) for 2.5 seconds by adding 20 µl of Nano-Glo® substrate (Promega Inc, WI, USA; diluted 1:40 in PBS). Individual unmixed populations of cells were treated identically and served as controls to determine basal luciferase activity to obtain normalized relative light units. The data were processed and plotted using GraphPad Prism 8 v8.4.3.

### Preparation of lentivirus particles carrying SARS-CoV-2 spike proteins for smFRET imaging

Lentiviruses carrying SARS-CoV-2 spikes were prepared similarly as previously described for HIV-1(*19-21, 54*). Two short peptides labeling tags (Q3: GQQQLG; A4: DSLDMLEM), either single or in pair-wise combinations were introduced into indicated positions in the S1 subunit on the plasmid pCMV-S (see Fig. S1). Viruses carrying 100 % single-tagged or double-tagged pCMV-S showed no noticeable defect in virus infectivity. The spike carrying one pair-wise combination of peptide tags (designated as pCMV-S Q3-1 A4-1) was used to make lentiviruses or S-MEN coronavirus like particles with SARS-CoV-2 spikes incorporated in the virus surface.

Lentivirus particles carrying SARS-CoV-2 spikes were produced by pseudotying an HIV-1 core with full-length spike proteins. To generate lentivirus particles that carry only one double tagged S on average among all the S trimers on the virus surface, plasmids encoding wildtype pCMV-S, pCMV-S Q3-1 A4-1, and pCMV delta R8.2 were transfected at a ratio of 20:1:21, respectively. Under these conditions, the vast majority of virus particles carry wildtype spikes. Among the small portion of virus particles containing tagged S, more than 95 % will carry one dually tagged protomer while the other two protomers remain wildtype. The same strategy has been deployed in our previous studies(*19-21, 54*). 293T cells were transfected using Fugene 6 with above indicated plasmids to express spikes on the HIV-1. When virus particles were generated that contained the disulfide bridge between S383C and D985C S, both plasmids, the plasmid encoding the spike protein, as well as plasmids encoding the dually tagged spike protein contained the S383C and D985C modification. To generate S-MEN particles, plasmids encoding wildtype pCMV-S, dual-tagged pCMV-S Q3-1 A4-1, pLVX-M (SARS-CoV-2 membrane protein expressing plasmid), an pLVX-E (SARS-CoV-2 envelope expressing plasmid, and pLVX-N (SARS-CoV-2 nucleocapsid expressing plasmid) were transfected at a ratio of 20:1:21:21:21. pLVX plasmids are gifts from Nevan Krogan, UCSF.

Lentivirus or S-MEN particles carrying dually-tagged S proteins were harvested 40 hour post-transfection, filtered through syringe filter with 0.45 mm pore size, and sedimented through a 15 % sucrose cushion at 25,000 rpm for 2 hours. The virus pellets were then re-suspended in 50 mM pH 7.5 HEPES buffer supplied with 10 mM MgCl_2_ and 10 mM CaCl_2_.

### Cryo-electron tomography

6 nm gold tracer was added to the concentrated S-decorated HIV-1 lentivirus and S-MEN particles viruses at 1:3 ratio, and 5□µl of the mixture was placed onto freshly glow discharged holey carbon grids for 1□min. Grids were blotted with filter paper, and rapidly frozen in liquid ethane using a homemade gravity-driven plunger apparatus.

Cryo-grids were imaged on a cryo-transmission electron microscope (Titan Krios, Thermo Fisher Scientific) that was operated at 300 kV, using a Gatan K3 direct electron detector in counting mode with a 20 eV energy slit. Tomographic tilt series between −51° and +51° were collected by using SerialEM(*55*) in a dose-symmetric scheme(*56*) with increments of 3°. The nominal magnification was 64,000 X, giving a pixel size of 1.346 Å on the specimen. The raw images were collected from single-axis tilt series with accumulative dose of ∼50 e− per Å^2^. The defocus was −3 μm and 8 frames were saved for each tilt angle.

Frames were motion-corrected using Motioncorr2(*57*) to generate drift-corrected stack files, which were aligned using gold fiducial makers by IMOD/etomo(*58*). Tomograms were reconstructed by weighted back projection and tomographic slices were visualized with IMOD.

### Fluorescently labeling spikes embedded on lentivirus particles

Virus particles were labeled through site-specifically enzymatic labeling, as previously described(*19, 21*). Transglutaminase transferred Cy3B(3S) from the cadaverine conjugate to the central glutamine residue of the Q3 (GQ<u>Q</u>QLG) tag in S1. The AcpS enzyme mediated the addition of the Cy5 derivative (LD650-CoA) to the serine residue of the A4 tag (D<u>S</u>LDMLEM). For these reactions to occur, Cy3B(3S)-cadaverine (0.5□ mM, Lumidyne Technologies), LD650-CoA (0.5 mM, Lumidyne Technologies), transglutaminase (0.65□ mM, Sigma Aldrich), and AcpS (5 mM, home-made) were added to the above suspensions of viruses. The labeling reaction mix was incubated at room temperature overnight. PEG2000-biotin (0.02 mg/ml, Avanti Polar Lipids) was added to the reaction mix and incubated for 30 min at room temperature with rotation. Free dyes and lipids were then purified away from labeled virus particles by ultracentrifugation for one hour at 4 °C at 40,000 rpm over a 6%-18% Optiprep (Sigma Aldrich) gradient. Purified virus particles were stored at −80 °C for future use in smFRET imaging.

### smFRET imaging data acquisition

All smFRET imaging data acquisition was performed on a home-built prism-based total internal reflection fluorescence (TIRF) microscope. Lentiviruses or S-MEN coronavirus-like particles carrying fluorescently-tagged spike protein, as well as lipid-biotin in the viral membranes, were immobilized on polyethylene glycol (PEG)-passivated, streptavidin-coated quartz slides for imaging. The evanescent field was generated at the interface between the quartz slide and the virus sample solution by prism-based total internal reflection, with laser excitation from a single-frequency 532-nm laser. Donor fluorophores labeled on viruses were directly excited by the generated evanescent field. Fluorescence signals from both donor and acceptor fluorophores were collected through a 1.27-NA 60 x water-immersion objective (Nikon). Collected signals were optically separated by passing through a 650 DCXR dichroic filter (Chroma) mounted on MultiCam LS image splitter (Cairn Research). Separated donor (ET590/50, Chroma) and acceptor fluorescent (ET690/50, Chroma) signals were simultaneously recorded on two synchronized ORCA-Flash4.0 V3 sCMOS cameras (Hamamatsu) at 25 frames per second for 80 seconds. Unless otherwise noted, all virus samples were imaged in pH 7.4, 50 mM Tris buffer with 50 mM NaCl, a cocktail of triplet-state quenchers, and an oxygen-scavenger system(*59*).

Where indicated, the conformational effects of different ligands/antibodies/plasma on SARS-CoV-2 spike were tested by pre-incubating fluorescently labeled viruses for 90 mins at room temperature before imaging in the continued presence of the ligands. In experiments with two different ligands presented, the second ligand (hACE2) was added to first-ligand-bound viruses and incubated similarly. The specific concentration of ligands incubated with viruses in smFRET imaging are as follows: 200 mg/ml hACE2 (human hACE2); 200 mg/ml dimeric hACE2, 50 mg/ml trypsin; 200 mg/ml CR3022; 200 mg/ml VHH72; 200 g/ml H4; 100 fold dilution of a SARS-CoV-2 positive patient plasma.

### smFRET data analysis

smFRET data analysis was performed using MATLAB (MathWorks)-based customized SPARTAN software package(*60*). For each labeled virus recorded over 80 seconds, the background signal at the single-molecule level was first identified based on the fluorophore bleaching point and subtracted. Donor and acceptor fluorescence intensity time-trajectory or traces were then extracted for each recorded virus and corrected for donor to acceptor crosstalk. The energy transfer efficiency (termed as FRET in graphs) from the donor to the neighboring acceptor was evaluated as FRET= I_A_/(γI_D_+I_A_), where I_D_ and I_A_ are the fluorescence intensities of donor and acceptor, respectively, and γ□ is the correlation coefficient compensating for the difference in quantum yields and detection efficiencies of donor and acceptor.

FRET efficiencies of each FRET pair report on the relative distance between the donor and the acceptor over time and ultimately reveal the conformational dynamics of host molecules (spikes on the virus in our case) in real-time. Viruses lacking donor or acceptor fluorophores, containing multiple labeled protomers, or containing more than one labeled spike on a single virus were automatically excluded from further analysis. FRET traces (FRET values as a function of real-time) were then manually selected if they displayed sufficient signal-to-noise (S/N) ratio and anti-correlated fluctuations in donor and acceptor fluorescence intensity between clearly defined FRET states, which are indicative of fully active molecules. These traces were compiled into FRET histograms. Based on visual inspection of traces that revealed direct observations of state-to-state transitions, FRET histograms were fitted into the sum of four Gaussian distributions using the least-squares fitting algorithm in Matlab. Each gaussian represents one FRET-indicated conformational state, and the area under each Gaussian curve estimates each state’s occupancy. The occupancy of each state was used to evaluate the difference in free energies between states *i* and *j*, according to Δ*G°*_*ij*_*=-k*_*B*_*Tln(P*_*i*_*/P*_*j*_), where *P*_*i*_ and *P*_*j*_ are the *i*th and *j*th state’s occupancy, respectively, *k*_*B*_ is the Boltzmann constant, and *T* is the temperature in kelvin.

The idealization of each FRET trace using a 4-state Hidden Markov Model was performed in the SPARTAN software package using a segmental *K*-means algorithm(*29*). The state-to-state transition tracing, indicative of the locations and the frequencies of transitions, was displayed in a transition density plot (termed as TDP). Dwell times (the time during which a molecule occupies one specific conformation before transitioning to any other conformations) distributions were compiled into survival probability plots and were fitted to the sum of two exponential distributions (y = *A*_1_ exp ^−*k*1*t*^ + *A*_2_ exp ^−*k*2*t*^). The transition rates were weighted from averaging the two rate constants by their amplitudes.

### Virus capture assay

The assay was modified from a previous published method(*39*). Briefly, pseudoviral particles were produced by transfecting 2×10^6^ HEK293T cells with pNL4.3 Luc R-E-(3.5 μg), plasmids(*38, 53*) encoding for SARS-CoV-2 Spike or SARS-CoV Spike (3.5 μg) protein and VSV-G (pSVCMV-IN-VSV-G, 1 μg) using the standard calcium phosphate protocol. Forty-eight hours later, supernatant-containing virion was collected and cell debris was removed through centrifugation (1,500 rpm for 10 min). To immobilize antibodies on ELISA plates, white MaxiSorp ELISA plates (Thermo Fisher Scientific) were incubated with 5 μg/ml of antibodies or 1:500 diluted plasma in 100 μl phosphate-buffered saline (PBS) overnight at 4 °C. Unbound antibodies or plasma were removed by washing the plates twice with PBS. Plates were subsequently blocked with 3% bovine serum albumin (BSA) in PBS for 1 hour at room temperature. After two washes with PBS, 200 μl of virus-containing supernatant was added to the wells. For sACE2 competition, 10 μM sACE2 was incubated with the supernatant at 37 °C for 1 hour before adding to the coated wells. After 4 to 6 hours incubation, supernatant was removed and the wells were washed with PBS 3 times. Viral capture by any given antibody was visualized by adding 10×10^4^ SARS-CoV-2-resistant Cf2Th cells (not shown) in full DMEM medium per well. Forty-eight hours post-infection, cells were lysed by the addition of 30 μl of passive lysis buffer (Promega) and three freeze-thaw cycles. An LB941 TriStar luminometer (Berthold Technologies) was used to measure the luciferase activity of each well after the addition of 100 μl of luciferin buffer (15 mM MgSO_4_, 15 mM KH_2_PO_4_ [pH 7.8], 1 mM ATP, and 1 mM dithiothreitol) and 50 μl of 1 mM D-luciferin potassium salt (Prolume).

### Flow cytometry analysis of cell-surface staining

Using the standard calcium phosphate method, 10 μg of Spike expressor and 2 μg of a green fluorescent protein (GFP) expressor (pIRES-GFP) was transfected into 2 × 10^6^ 293T cells. At 48 hours post transfection, 293T cells were stained with plasma from SARS-CoV-2-infected or uninfected individuals (1:250 dilution). The percentage of transfected cells (GFP+ cells) was determined by gating the living cell population based on the basis of viability dye staining (Aqua Vivid, Invitrogen). Samples were acquired on a LSRII cytometer (BD Biosciences, Mississauga, ON, Canada) and data analysis was performed using FlowJo vX.0.7 (Tree Star, Ashland, OR, USA). Alternatively, convalescent plasma was incubated with 20 μg/mL of SARS-CoV-2 RBD prior cell-surface staining in order to compete for RBD-specific antibodies.

### ELISA (enzyme-linked immunosorbent assay)

The SARS-CoV-2 RBD ELISA assay used was recently described(*38, 53*). Briefly, recombinant SARS-CoV-2 S RBD proteins (2.5 μg/ml), or bovine serum albumin (BSA) (2.5 μg/ml) as a negative control, were prepared in PBS and were adsorbed to plates (MaxiSorp; Nunc) overnight at 4 °C. Coated wells were subsequently blocked with blocking buffer (Tris-buffered saline [TBS] containing 0.1% Tween20 and 2% BSA) for 1 hour at room temperature. Wells were then washed four times with washing buffer (Tris-buffered saline [TBS] containing 0.1% Tween20). CR3022 mAb (50 ng/ml) or serial dilutions of plasma from SARS-CoV-2-infected or uninfected donors (1/100; 1/250; 1/500; 1/1000; 1/2000; 1/4000) were prepared in a diluted solution of blocking buffer (0.1 % BSA) and incubated with the RBD-coated wells for 90 minutes at room temperature. Plates were washed four times with washing buffer followed by incubation with anti-IgG secondary Abs (diluted in a diluted solution of blocking buffer [0.4% BSA]) for 1hour at room temperature, followed by four washes. HRP enzyme activity was determined after the addition of a 1:1 mix of Western Lightning oxidizing and luminol reagents (Perkin Elmer Life Sciences). Light emission was measured with a LB941 TriStar luminometer (Berthold Technologies). Signal obtained with BSA was subtracted for each plasma and was then normalized to the signal obtained with CR3022 mAb present in each plate.

---

**Fig. S1.**
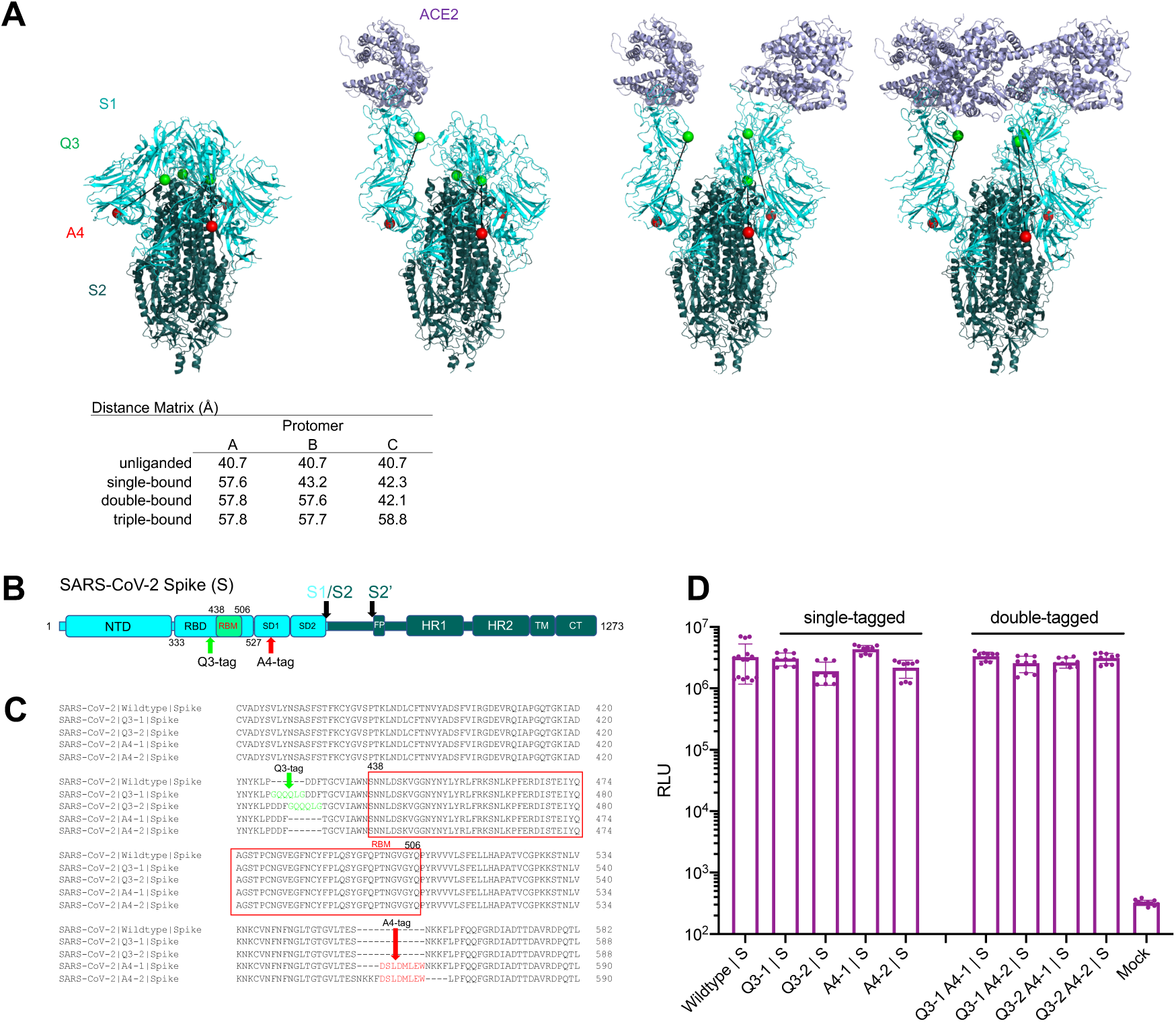
Single-tagged and double-tagged spike proteins for smFRET imaging retain infectivity. **(A)** CryoEM structures of a soluble trimeric SARS-CoV-2 S, with GSAS and PP mutations and the T4 phage fibritin trimerization domain(*17*). ligand-free (PDB: 6VXX) and recognizing single, double and triple ACE2 receptors molecules(*18*), with matrix showing distances between Q3-1 and A4-1 insertion sites on S1. **(B)** Domain organization of full length (1273 amino acids) wildtype SARS-CoV-2 spike protein (S1+S2) colored by domain. The insertion sites of labeling tags Q3 and A4 for the donor and acceptor fluorophores are indicated. S1/S2, S2’, protease cleavage sites; NTD, N-terminal domain; RBD, receptor-binding domain; RBM, receptor binding motif; SD1, subunit domain 1; SD2, subunit domain 2; FP, fusion peptide; HR1/HR2, heptad repeat 1/heptad repeat 2; TM, transmembrane domain; CT, cytoplasmic tail. **(C)** Peptide insertion sites into RBD and SD1 before and after RBM to avoid interfering with receptor binding. Q3 tag was inserted at two positions (Q3-1 and Q3-2) on RBD, separately. A4 peptide tag was inserted at two sites (A4-1 and A4-2) on SD1. **(D)** Infectivity of HIV-1 lentivirus carrying single-tagged and double-tagged spike proteins as determined on Huh 7.5 cells. Infectivity (mean ± s.d.) was measured from three independent experiments in triplicates. RLU, relative light units.

**Fig. S2.**
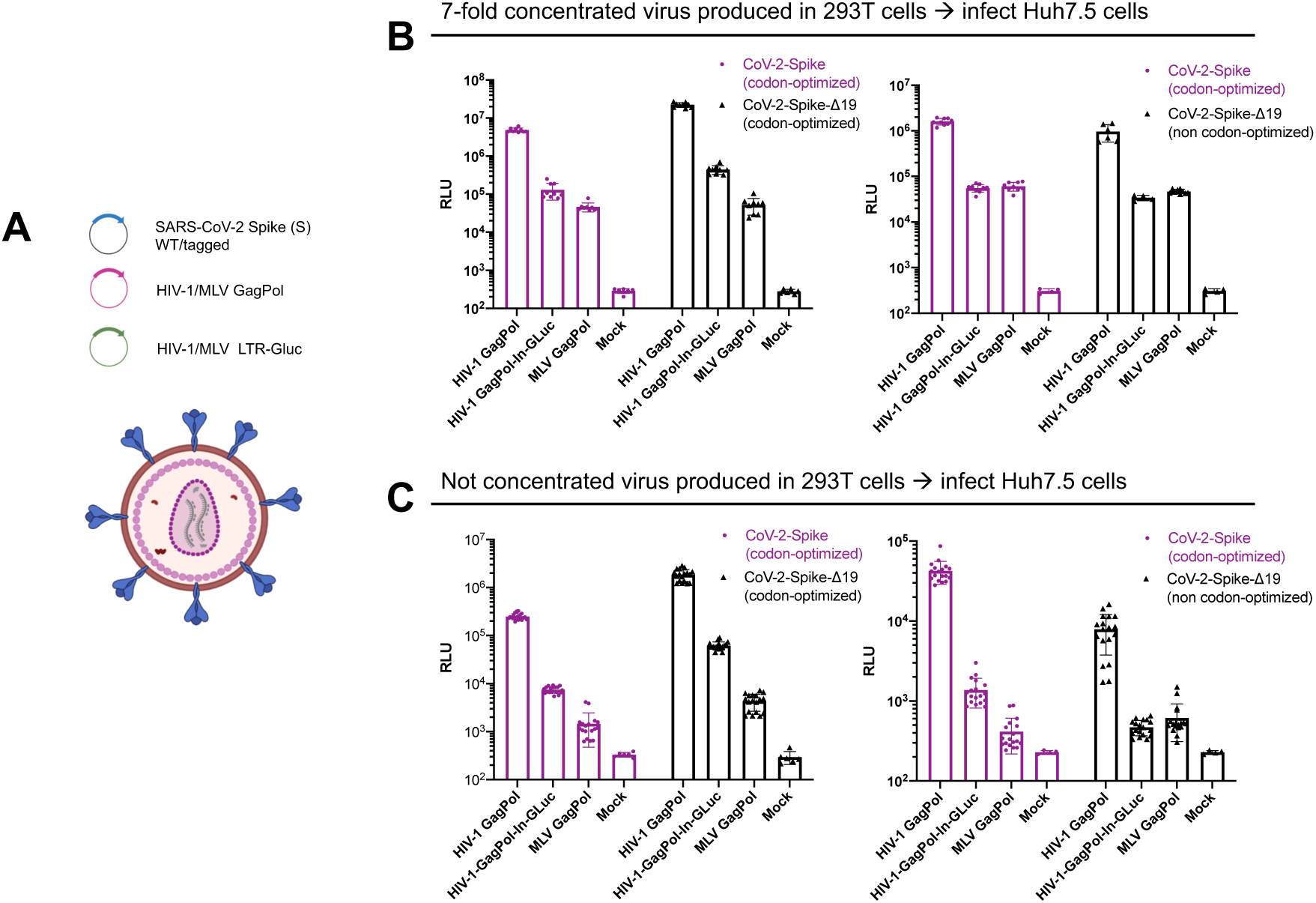
Establishment of infectivity assays using lentivirus and retrovirus particles carrying SARS-CoV-2 spike proteins. **(A)** HIV-1 or MLV single-round viruses used for infectivity and neutralization assays were generated by transfecting HEK293T cells with plasmids expressing respective GagPol, LTR-intron Gaussia luciferase Gluc(*50, 51*) or plasmid encoding HIV-1 envelope-deleted genome encoding intron Gluc together with codon-optimized, non-codon-optimized or Δ19 (spike with 19 amino acids deleted from C-terminal) versions of SARS-CoV-2 spike(*52*). **(B)** Graphs comparing infectivity of indicated viruses titered on Huh 7.5 measured as a function of Gaussia luciferase activity (relative light units, RLU; mean ± s.d.) 48 h after infection from 7-fold concentrated or neat culture supernatants (**C**). Viral infectivity assays were carried out three independent times in triplicates.

**Fig. S3.**
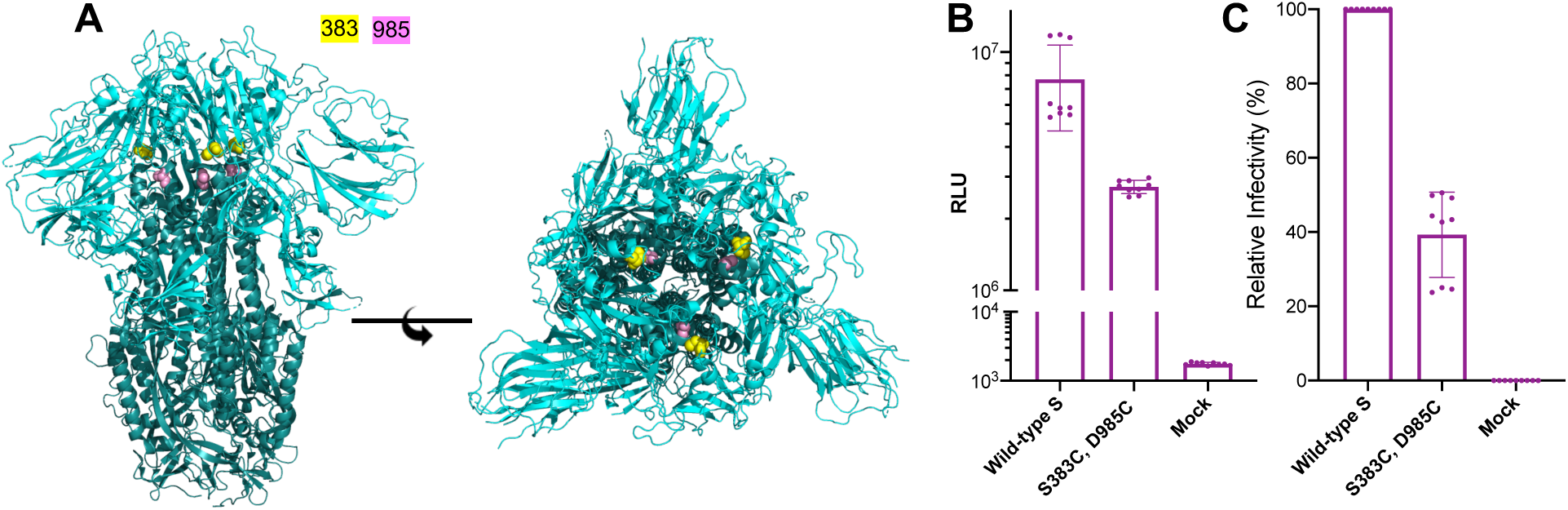
Relative infectivity of S containing S383C, D985C modifications. **(A)** Structure (PDB: 6ZOY) of disulfide (S383C, D985C)-stabilized spike in all ‘down’ state (left panel: side view; right panel: top view; 383 site in yellow spheres; 985 site in pink spheres). **(B, C)** Infectivity of spike mutant S383C, D985C in comparison to wildtype, represented as absolute RLU (**B**) and normalized (**C**) to wild-type (%). Infectivity (mean ± s.d.) was repeated three times in triplicates. RLU, relative light units.

**Fig. S4.**
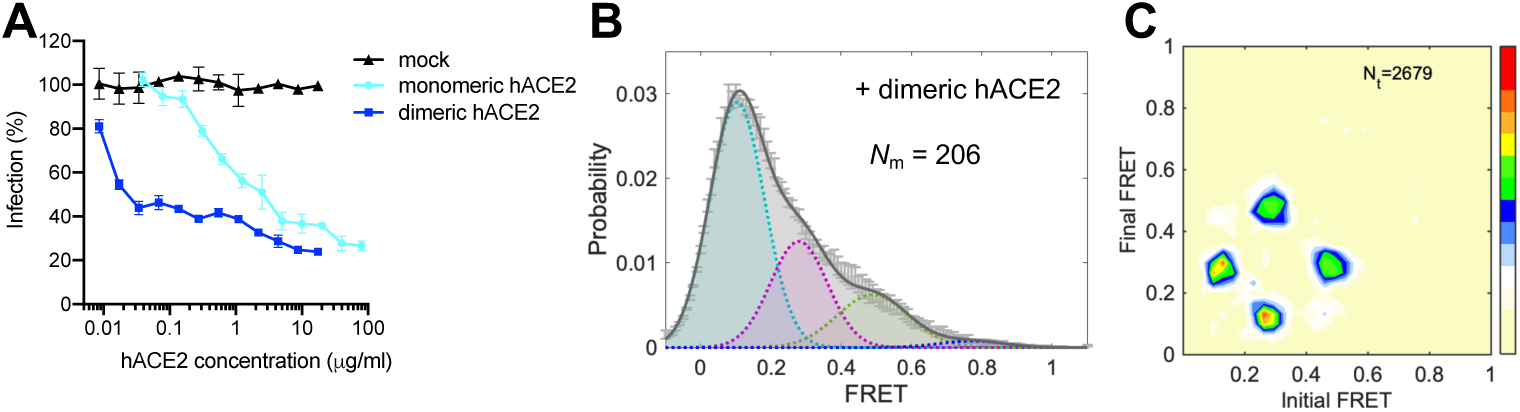
Dimeric hACE2 activates spike protein by stabilizing the low-FRET conformation. **(A)** Dimeric hACE2 is more potent than monomeric hACE2. Infectivity of lentivirus particles carrying SARS-CoV-2 Spike in presence of indicated amounts of soluble monomeric or dimeric hACE2 protein using 293T cells stably expressing hACE2 as target cells. Experiment was performed in triplicates and represented as mean ± s.d. **(B, C)** The FRET histogram (**B**) and the corresponding transition density plot (**C**) of spikes on lentivirus particles in the presence of 200 μg/ml dimeric hACE2. 206 (*N*_m_) individual dynamic traces were compiled into a conformation-population FRET histogram (gray lines, **B**) and fitted into a 4-state Gaussian distribution (solid black, **B**). Compared to monomeric hACE2, dimeric hACE2 further reduces the occupancy of the very high-FRET and facilitates the stabilization in the low-FRET receptor-stabilized state (**C**). FRET histograms represent mean ± s.e.m., determined from three randomly-assigned populations of all FRET traces. Evaluated state occupancies see Table S1.

**Fig. S5.**
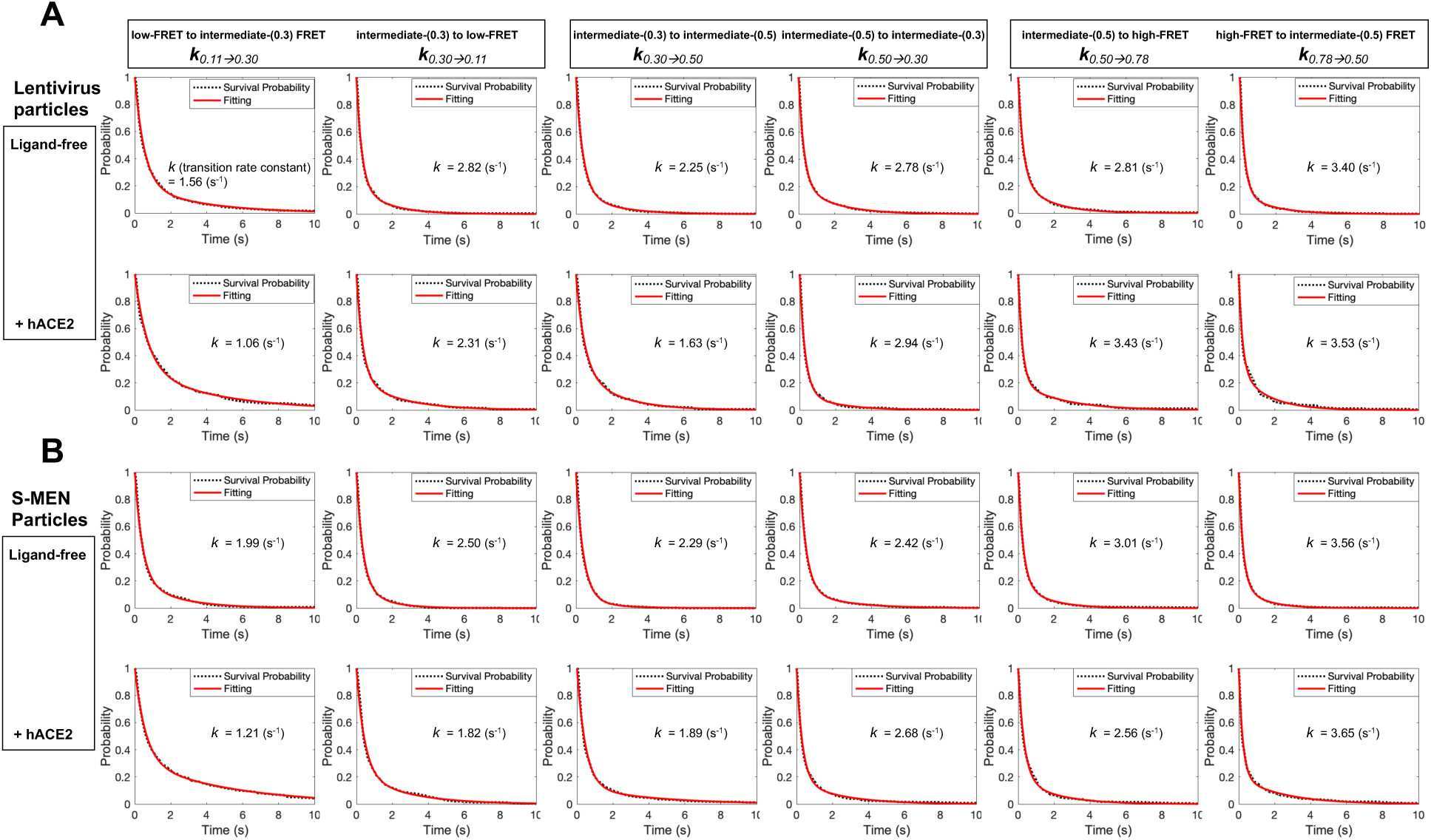
Kinetic analysis of spike proteins, and modulation by hACE2. **(A, B)** Survival probability plots of conformational states of spikes on lentivirus particles (**A**) and on S-MEN (**B**) in the absence (ligand-free, top row) and the presence of 200 μg/ml hACE2 (bottom row). The survival probability plot is a function of time-duration of spikes dwelling on specific conformations before state-to-state transitions occur. Transition rates were derived from double exponential-fitting of survival probability plots and summarized in **Table S2**.

**Fig. S6.**
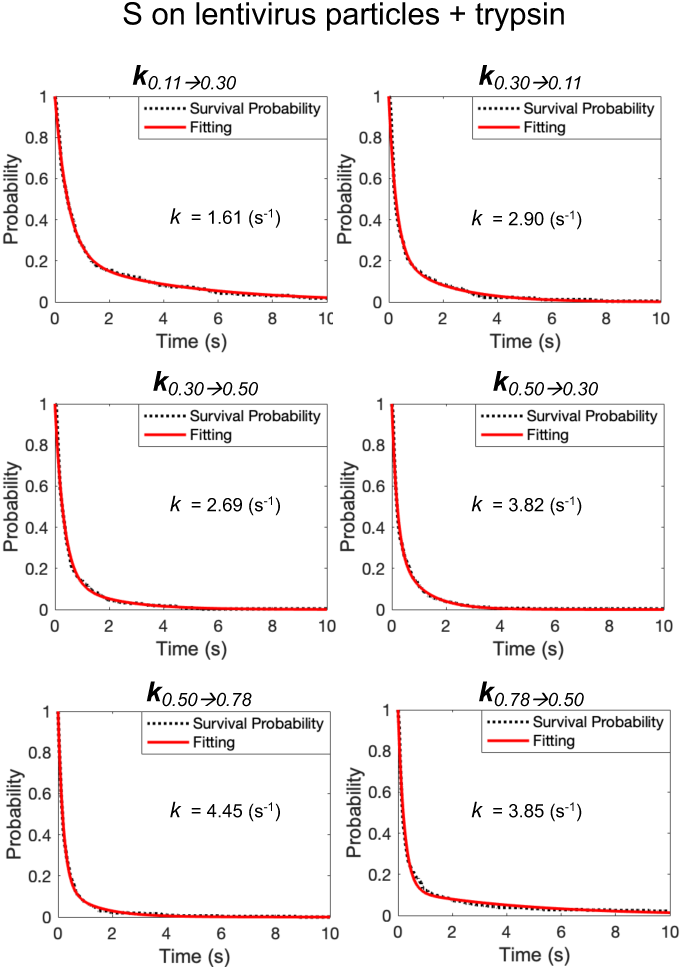
The effect of trypsin on conformational dynamics of SARS CoV-2 spike on lentivirus particles. Survival probability plots of spike-conformations in the presence of 50 ug/ml trypsin give rise to the calculation of transition rates among different conformations of spikes. Transition rates are summarized in **Table S2**.

**Fig. S7.**
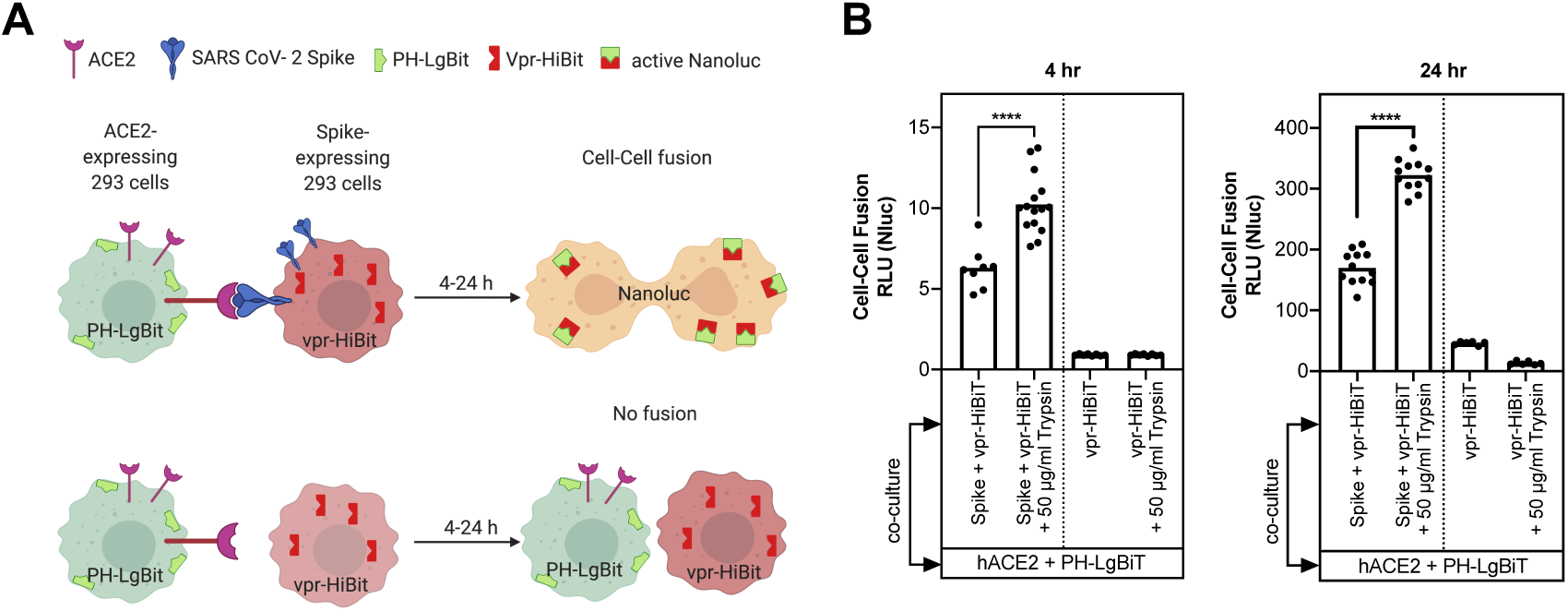
Effect of trypsin treatment on hACE2-SARS-CoV-2-mediated cell-cell fusion. **(A)** Assay design to monitor cell-cell fusion using the HiBit and LgBiT split nanoluc system(*47*). HEK293 cells transiently expressing Vpr-HiBit with or without SARS-CoV-2 spike were co-cultured with HEK293 cells transiently expressing LgBiT tagged to PH domain of human phospholipase Cd at the N-terminus together with hACE2. SARS-CoV-2 spike-hACE2-dependent cell-cell fusion was determined by monitoring reconstituted nanoluc activity at 4 and 24 h post co-culture. **(B)** Normalized relative luciferase units (RLU; bars denote mean; dots, individual data points of replicates) measured at 4 h and 24 h post co-culture to quantify cell-cell fusion in stated cell populations. Indicated cell populations were either left alone or treated with 50 µg/ml trypsin for 10 min at 37 °C before co-culture. Reconstituted NanoLuc activities were normalized to luciferase activity detected in cell populations without co-culture. p values were derived from unpaired t-test; **** denote p < 0.0001

**Fig. S8.**
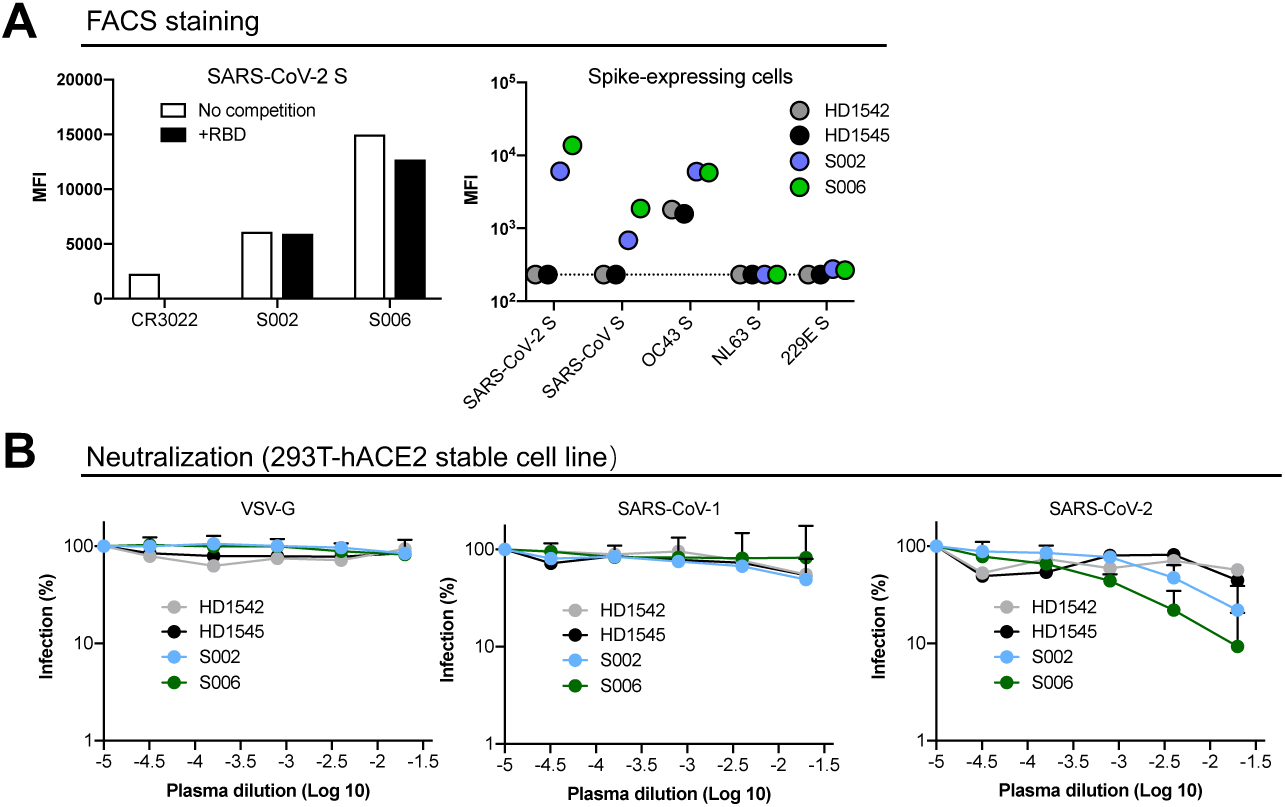
Convalescent plasma of SARS-CoV-2 positive patients S006 and S002 bind to S and neutralize SARS-CoV-2. **(A)** FACS staining of SARS-CoV-2 S-expressing cells shows that both S006 and S002 bind to S. **(B)** VSV-G (left), SARS-CoV-1 (middle), SARS-CoV-2 (right) lentivirus neutralization curves of plasma S006, S002, HD1542 and HD1545. Neutralization curves represent mean ± s.d. from three replicates.

**Fig. S9.**
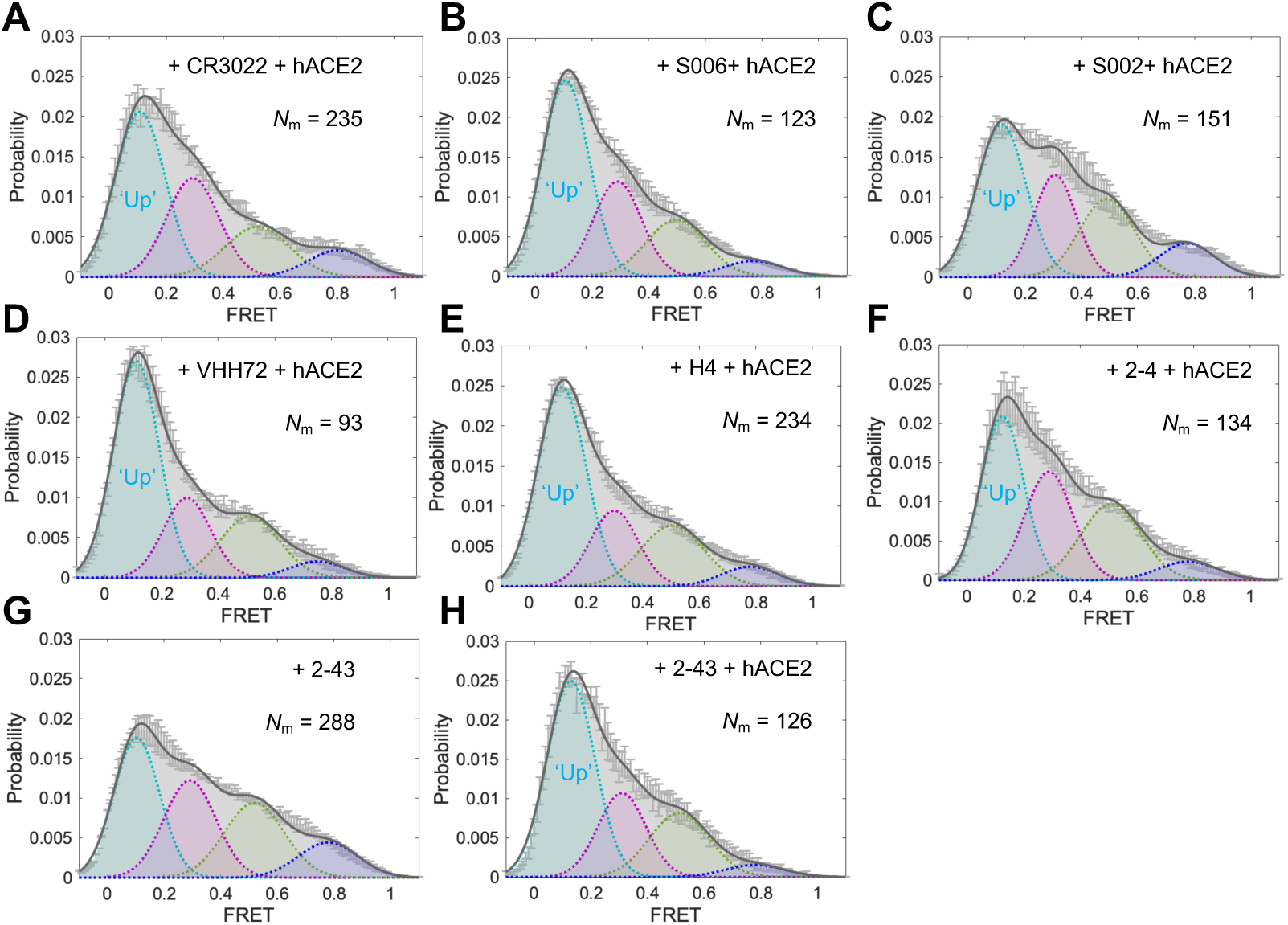
hACE2 mediate the antibodies-equilibrated conformational landscape towards down-stream conformations. **(A–F)** FRET histograms of SARS-CoV-2 S on the lentivirus in the presence of both 200 μg/ml of indicated antibody and 200 μg/ml hACE2, or 1:100 dilution of indicated plasma and 200 μg/ml hACE2. (**G, H)** Experiments as in **A–F**, in the absence and presence of 200 μg/ml hACE2 in addition to 200 μg/ml of 2-43 antibody. FRET histograms represent mean ± s.e.m., determined from three randomly assigned populations of all FRET traces. Evaluated state occupancies see Table S1.

**Table S1.** Relative state-occupancy and fitting parameters in each of four FRET-defined conformational states of SARS-CoV-2 spike protein on the surface of virus particles. The FRET efficiency histograms were fitted into the sum of four Gaussian distributions (μ, the mean or expectation of the Gaussian distribution; σ, s.d. of the Gaussian distribution) for each conformational state. Parameters were based upon the observation of original FRET efficiency data and were further determined using hidden Markov modelling. Relative conformational state-occupancy of SARS-CoV-2 spike protein on viral particles are presented as mean ± s.e.m., determined from three independent measurements. R-squared values were evaluated to indicate the goodness of fit.

| SARS-CoV-2 spikes on lentivirus particles | Curve fitting R-squared | low-FRET | intermediate-(0.3) FRET | intermediate-(0.5) FRET | high-FRET |
| --- | --- | --- | --- | --- | --- |
| | | $\mu$: 0.11±0.01 | $\mu$: 0.30±0.01 | $\mu$: 0.50±0.02 | $\mu$: 0.78±0.03 |
| | | $\sigma$: 0.08±0.01 | $\sigma$: 0.08±0.01 | $\sigma$: 0.10±0.01 | $\sigma$: 0.10±0.01 |
| Ligand-free | 0.9921 | 22% ± 7% | 26% ± 11% | 38% ± 12% | 14% ± 8% |
| + hACE2 | 0.9966 | 48% ± 10% | 29% ± 11% | 17% ± 12% | 6% ± 3% |
| + dimeric hACE2 | 0.9968 | 58% ± 8% | 25% ± 12% | 16% ± 11% | 2% ± 2% |
| + trypsin | 0.9808 | 28% ± 9% | 27% ± 9% | 28% ± 9% | 17% ± 5% |
| + trypsin + hACE2 | 0.9882 | 51% ± 9% | 24% ± 11% | 19% ± 10% | 5% ± 8% |
| + CR3022 | 0.9952 | 45% ± 3% | 24% ± 10% | 21% ± 9% | 10% ± 7% |
| + CR3022 + hACE2 | 0.9926 | 44% ± 8% | 28% ± 13% | 18% ± 12% | 10% ± 8% |
| + VHH72 | 0.9915 | 46% ± 6% | 25% ± 12% | 20% ± 12% | 9% ± 4% |
| + VHH72 + hACE2 | 0.9949 | 55% ± 7% | 20% ± 9% | 20% ± 10% | 5% ± 7% |
| + H4 | 0.9947 | 51% ± 9% | 22% ± 12% | 22% ± 12% | 5% ± 6% |
| + H4 + hACE2 | 0.9966 | 53% ± 8% | 20% ± 12% | 21% ± 13% | 6% ± 4% |
| + S002 | 0.9958 | 23% ± 5% | 20% ± 12% | 45% ± 12% | 12% ± 6% |
| + S002 + hACE2 | 0.9904 | 41% ± 7% | 24% ± 12% | 24% ± 12% | 11% ± 4% |
| + S006 | 0.9930 | 54% ± 11% | 24% ± 12% | 16% ± 10% | 6% ± 3% |
| + S006 + hACE2 | 0.9955 | 51% ± 10% | 25% ± 12% | 19% ± 13% | 5% ± 5% |
| + 2-4 | 0.9907 | 25% ± 10% | 20% ± 13% | 45% ± 11% | 10% ± 5% |
| + 2-4 + hACE2 | 0.9959 | 39% ± 10% | 29% ± 13% | 26% ± 12% | 6% ± 5% |
| + 2-43 | 0.9925 | 35% ± 3% | 28% ± 11% | 25% ± 8% | 12% ± 5% |
| + 2-43 + hACE2 | 0.9906 | 52% ± 9% | 22% ± 13% | 22% ± 13% | 4% ± 5% |
| S383C, D985C ligand-free | 0.9933 | 20% ± 5% | 18% ± 11% | 45% ± 12% | 17% ± 7% |

| SARS-CoV-2 S-MEN | Curve fitting R-squared | low-FRET | intermediate-(0.3) FRET | intermediate-(0.5) FRET | high-FRET |
| --- | --- | --- | --- | --- | --- |
| | | $\mu$: 0.11±0.01 | $\mu$: 0.30±0.01 | $\mu$: 0.50±0.02 | $\mu$: 0.78±0.03 |
| | | $\sigma$: 0.08±0.01 | $\sigma$: 0.08±0.01 | $\sigma$: 0.10±0.01 | $\sigma$: 0.10±0.01 |
| Ligand-free | 0.9886 | 26% ± 6% | 23% ± 11% | 39% ± 12% | 12% ± 7% |
| + hACE2 | 0.9977 | 49% ± 8% | 24% ± 11% | 20% ± 11% | 7% ± 6% |

**Table S2.** Transition rates between observed conformational states of SARS-CoV-2 spike on virus particles. The survival probability plots (Figs. S5 and S6) were derived from distributions of dwell times for each state-to-state transitions determined through Hidden Markov Modeling (HMM). Then plots were fitted by double-exponential distributions: *y(t)* = *A*_*1*_ exp ^− *k*1*t*^ + *A*_2_ exp ^−*k*2*t*^), where *y(t)* is the probability and *t* is the dwell time. The presented rates were the weighted average of two rates derived from double-exponential decays. Rates were finally presented in the table as (*weighted average* +/- *95% confidence intervals*).

| **SAR-CoV-2 Spike**<br>**on virus particles** | $k_{0.11 \rightarrow 0.30} (s^{-1})$ | $k_{0.30 \rightarrow 0.11} (s^{-1})$ | $k_{0.30 \rightarrow 0.50} (s^{-1})$ | $k_{0.50 \rightarrow 0.30} (s^{-1})$ | $k_{0.50 \rightarrow 0.78} (s^{-1})$ | $k_{0.78 \rightarrow 0.50} (s^{-1})$ |
| --- | --- | --- | --- | --- | --- | --- |
| **Retroviral**<br>**pseudo-particles** |  |  |  |  |  |  |
| Ligand-free | 1.56 +/- 0.01 | 2.82 +/- 0.03 | 2.25 +/- 0.03 | 2.78 +/- 0.02 | 2.81 +/- 0.02 | 3.40 +/- 0.03 |
| + hACE2 | 1.06 +/- 0.01 | 2.31 +/- 0.03 | 1.63 +/- 0.03 | 2.94 +/- 0.03 | 3.43 +/- 0.04 | 3.53 +/- 0.07 |
| + Trypsin | 1.61 +/- 0.01 | 2.90 +/- 0.05 | 2.69 +/- 0.04 | 3.82 +/- 0.07 | 4.45 +/- 0.04 | 3.85 +/- 0.04 |
| **S-MEN particles** |  |  |  |  |  |  |
| Ligand-free | 1.99 +/- 0.02 | 2.50 +/- 0.04 | 2.29 +/- 0.04 | 2.42 +/- 0.01 | 3.01 +/- 0.03 | 3.56 +/- 0.03 |
| + hACE2 | 1.21 +/- 0.01 | 1.82 +/- 0.02 | 1.89 +/- 0.02 | 2.68 +/- 0.03 | 2.56 +/- 0.03 | 3.65 +/- 0.04 |

